# CRISPR activation and interference screens in primary human T cells decode cytokine regulation

**DOI:** 10.1101/2021.05.11.443701

**Authors:** Ralf Schmidt, Zachary Steinhart, Madeline Layeghi, Jacob W. Freimer, Vinh Q. Nguyen, Franziska Blaeschke, Alexander Marson

## Abstract

The pathways that regulate cytokine responses in T cells are disrupted in autoimmunity, immune deficiencies, and cancer, and include immunotherapy targets. Systematic discovery of cytokine regulators requires both loss-of-function and gain-of-function studies, which have been challenging in primary human cells. We now have accomplished genome-wide pooled CRISPR activation (CRISPRa) and CRISPR interference (CRISPRi) screens in primary human T cells to map gene networks controlling Interleukin-2 and Interferon-γ production. Arrayed CRISPRa confirmed key hits and enabled multiplexed T cell secretome characterization, revealing reshaped cytokine responses driven by individual regulators. CRISPRa uncovered genes not canonically expressed in T cells, including the transcription factor *FOXQ1,* whose overexpression promoted the expression of most cytokines, while selectively dampening T helper 2 (Th2) cytokines. Paired CRISPRa and CRISPRi screens reveal signaling components that tune critical immune cell functions, which could inform design of future immunotherapies.

## Main Text

Cytokines produced by T cells upon stimulation are a critical part of the adaptive immune response. The set of cytokines produced by specific T cell subsets are tightly regulated to ensure effective immune reactions while preventing excessive inflammation. Dysregulated T cell cytokine responses are linked to a multitude of pathological conditions including autoimmunity, immunodeficiency, and immune evasion in cancer (*1*–*4*). Interleukin-2 (IL-2) is a cytokine predominantly secreted by CD4^+^ T cells, is a major driver of T cell expansion during adaptive immune responses (*5*), and has been developed as an immunotherapy for cancer and autoimmunity at different doses (*6*). Interferon-γ (IFN-γ) is a cytokine secreted by both CD4^+^ and CD8^+^ T cells that promotes a type I immune response, resulting in the induction of T helper 1 (Th1) CD4^+^ T cells, cytotoxic CD8^+^ T cells and IgG2a class switched B cells (*4*). Furthermore, IFN-γ plays a direct role in tumor killing, is correlated with immunotherapy response (*7*), and resistance to IFN-γ is a mechanism of immune escape in malignant cells (*8*, *9*). Much of our current understanding of the pathways leading to cytokine production in humans originated from studies in transformed T cell lines, which often are not representative of primary human cell biology (*10*–*12*). Comprehensive understanding of pathways that control cytokine production in primary human T cells is necessary to facilitate the development of next generation immunotherapies.

Unbiased forward genetic approaches can uncover the components of regulatory networks systematically. Genome-wide CRISPR knockout screens have been completed using primary mouse immune cells from Cas9 expressing transgenic mice (*13*–*15*), including a screen for regulators of innate cytokine response in dendritic cells (*13*), however CRISPR screens in primary human cells have been more challenging due to the requirement of efficient Cas9 delivery. We recently developed the SLICE (sgRNA lentiviral infection with Cas9 protein electroporation) platform for pooled genome-wide CRISPR knockout screens in primary human T cells (*16*). While CRISPR knockout identifies genes essential for a specific phenotype, it misses regulators that are active outside of the screening parameters (*17*). Scalable gain-of-function screening approaches, such as CRISPRa, have the potential to overcome this limitation, and furthermore may discover genes that act as regulatory “bottlenecks.” However, unlike CRISPR knockout, CRISPRa requires sustained expression of an activator-linked endonuclease-dead Cas9 (dCas9), which cannot be readily achieved with protein or mRNA electroporation. CRISPRa has been established in immortalized human cell lines through viral transduction and transgene integration, but lentiviral delivery of Cas9 gene constructs have been challenging in primary human T cells, limiting the use of CRISPRa to small scale experiments in primary cells to date (*18*, *19*).

To overcome lentiviral delivery challenges and enable scalable CRISPRa in primary human T cells, we developed an optimized high-titer lentiviral production protocol with a minimal dCas9-VP64 vector (pZR112), allowing for transduction efficiencies up to 80% (fig. S1). We confirmed CRISPRa efficacy with a panel of established sgRNAs targeting the transcriptional start sites (TSS) of genes encoding surface receptors. A second-generation CRISPRa synergistic activation mediator (SAM) system (*20*, *21*), consisting of dCas9-VP64 and aptamer-based recruitment of P65 and HSF1 transcriptional activators, induced robust increases in target surface marker expression (fig. S2). Next, we scaled up our CRISPRa platform to perform pooled genome-wide CRISPRa screens using the Calabrese library, targeting the TSS’s of >18,800 protein coding genes (*20*). We used fluorescent activated cell sorting (FACS) to sort for IL-2 producing CD4^+^ T cells and IFN-γ producing CD8^+^ T cells from two donors into high and low bins, respectively (Fig. 1A and fig. S3A-C). Subsequent extraction of genomic DNA and sgRNA quantification confirmed sgRNAs targeting the genes encoding IL-2 (*IL2)* and IFN-γ (*IFNG*) were strongly enriched in the respective positive bins (Fig. 1B). In contrast, non-targeting control sgRNAs were not enriched in high or low sorting bins (Fig. 1B). Both CRISPRa screens were highly reproducible with strong donor-to-donor correlation (Fig. 1C-D and fig. S3D-E). Gene level statistical analysis of the IL-2 and IFN-γ CRISPRa screens revealed 444 and 471 hits, respectively, including 171 shared hits (Fig. 1E, fig. S3F-G, and tables S1, S2).

**Figure 1.**
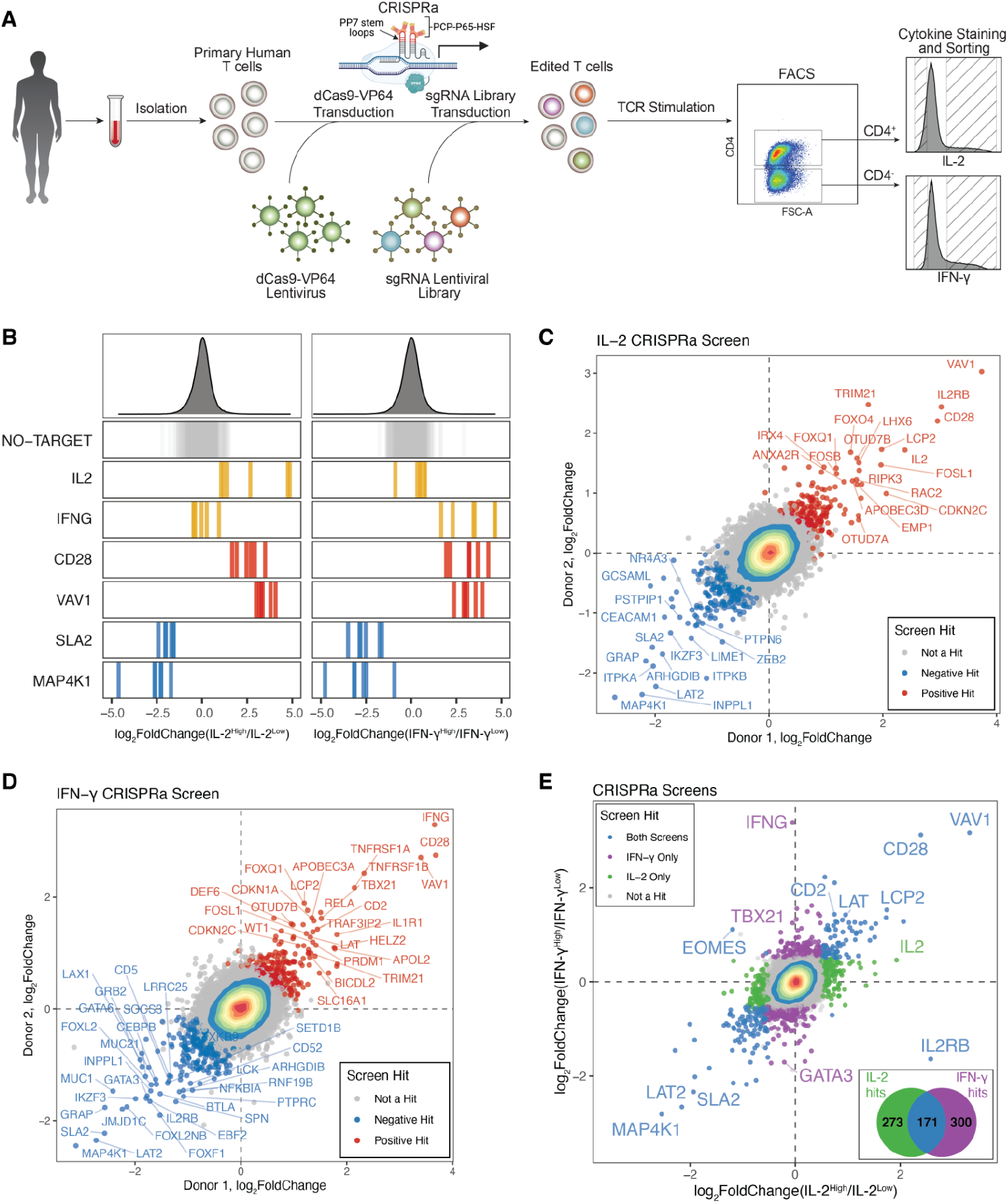
Genome-wide CRISPRa screens for cytokine production in primary human T cells. **(A)** Schematic of CRISPRa screens. **(B)** sgRNA log_2_ fold changes for genes of interest in IL-2 (left) and IFN-γ (right) screens. Bars represent the mean log_2_ fold change for each sgRNA across two donors. Density plots above represent the distribution of all sgRNAs. **(C-D)** Scatter plots of median sgRNA log_2_ fold change (high/low sorting bins) for each gene, comparing screens in two donors, for IL-2 (C) and IFN-γ screens (D). **(E)** Comparison of gene log_2_ fold change (median sgRNA, mean of two donors) in IL-2 and IFN-γ screens. Sub-panel Venn-diagram shows the number of screen unique and shared hits.

We discovered genes that could be tuned by CRIPSRa to shape cytokine responses. These included well known components of the TCR signaling pathway, and T cell transcription factors. Activation of *TBX21* (encoding T-bet), the lineage defining transcription factor of Type I immunity promoting Th1 CD4^+^ T cells, specifically enhanced IFN-γ production (Fig.1E), the signature Th1 cytokine (*22*, *23*). Conversely, sgRNAs activating *GATA3*, which promotes Th2 differentiation by antagonizing T-bet (*22*, *24*), were enriched in the IFN-γ^Low^ bin (Fig. 1E). Overexpression of genes encoding members of the proximal TCR/co-stimulatory signal transduction complex, such as *VAV1*, *CD28, LCP2* (encoding SLP-76), and *LAT (25, 26)* reinforced general T cell activation, supported by their enrichment in both cytokine high bins. Consistent with these observations, known negative regulators of proximal TCR signaling components, *MAP4K1 and SLA2* were depleted in these bins (Fig. 1B, E). We conclude that CRISPRa is able to identify critical bottlenecks in signals leading to cytokine responses that can be overcome by increasing levels of specific signaling components.

CRISPRa screens were effective in identifying limiting factors in cytokine response, but we recognized they could miss essential components that would only be identified through loss-of-function studies. We therefore performed reciprocal genome-wide CRISPRi screens, adapting the lentiviral protocols optimized for CRISPRa and employing the Dolcetto library (*27*) (Fig 2A, fig. S4A-B, and tables S1, S2). We validated the quality of CRISPRi screens by confirming that sgRNAs targeting gold-standard essential genes (*28*) dropped out of the population during T cell expansion (fig. S5). CRISPRi cytokine screens were also highly reproducible across donors, with the IL-2 and IFN-γ screens identifying 226 and 203 gene hits, respectively, including 92 shared hits (fig. S4C-H). We next compared and contrasted regulators identified by CRISPRi and CRISPRa. Despite CRISPRa identifying many well-established proximal regulators of TCR signaling, it did not identify some key known positive regulators, such as the CD3 surface complex, encoded by *CD3D, CD3E,* and *CD3G* genes, whereas CRISPRi identified all three genes as top ranked hits (fig. 2B and fig. S6A). This suggests that although CD3 complex genes are required for signaling, they are likely not bottlenecks in the pathway so signaling cannot be enhanced by their overexpression.

**Figure 2.**
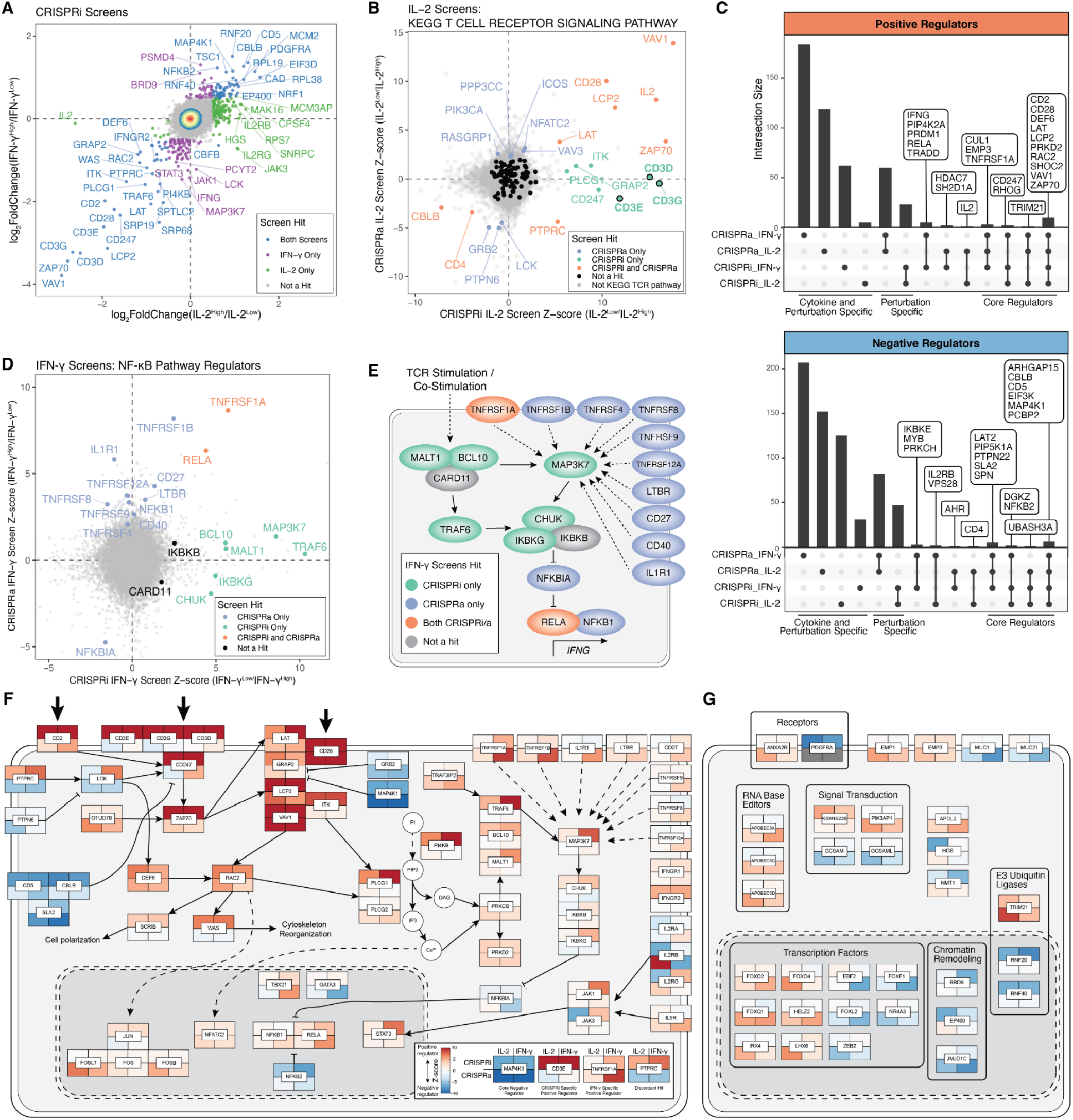
Integrated CRISPRa and CRISPRi screens map the genetic circuits underlying T cell cytokine response in high resolution. **(A)** Comparison of gene log_2_ fold change (median sgRNA) in IL-2 and IFN-γ CRISPRi screens. **(B)** Comparison IL-2 CRISPRi and CRISPRa screens with genes belonging to the T cell receptor signaling pathway (KEGG pathways) indicated in colors other than grey. Genes meeting hit criteria are labeled. CD3 surface complex genes are indicated by black border and bolded labels. Scales represent z-score of log_2_ fold change, with positive regulators of IL-2 production having positive z-scores. **(C)** Unique and common positive and negative regulators identified across CRISPRa and CRISPRi screens. For CRISPRa, positive regulators are defined as having a positive log_2_ fold change (high/low bins), and for CRISPRi a negative log_2_ fold change (high/low). **(D)** Comparison IFN-γ CRISPRi and CRISPRa screens with manually selected NF-κB pathway regulators labeled. All other genes are shown in grey. **(E)** Map of NF-κB pathway regulators labeled in (D). **(F)** Map of screen hits with previous evidence of defined function in T cell stimulation and costimulation signal transduction pathways. Tiles represent proteins encoded by indicated genes, with the caveat that due to space constraints, subcellular localization is inaccurate, as many of the components shown in the cytoplasm occur at the plasma membrane. Tiles are colored according to log2 fold change z-score as shown in the sub-panel, with examples of different hits. Large arrows at the top represent stimulation/co-stimulation sources by Immunocult. **(G)** Map of top screen hits with previously undescribed or poorly described function in T cells in the same format as (F).

To gain insights into functional pathways enriched across CRISPRi and CRISPRa screens, we completed gene set enrichment analysis of KEGG pathways, identifying multiple immune related pathways as enriched across four screens (fig. S6B). Given similar pathway enrichments across screens, we asked if this overlap occurs at the gene level. Integrative analysis of CRISPRa and CRISPRi screen hits found that a handful of genes were identified across all screens (e.g. *ZAP70* as a positive regulator and *CBLB* as a negative regulator), representing core regulators of T cell cytokine response to stimulation, whereas the majority of hits are either cytokine (IL-2 or IFN-γ) or perturbation (activation or interference) specific (Fig. 2C). Notably, CRISRPa and CRISPRi had discordant effects on cytokine production for 14 genes (e.g. *PTPRC*), suggesting that for some genes activation and interference both impair optimal levels (fig. S6C). Taken together, CRISPRi and CRISPRa hits reveal core and context-specific regulators of cytokine production.

The power of combinatorial activation and interference screening was exemplified by hits encoding members of the tumor necrosis factor receptor superfamily (TNFRSF) and the nuclear factor kappa B (NF-κB) signaling pathway. CRISPRi screens identified a key circuit of TCR stimulation signaling through MALT1, BCL10, TRAF6, and TAK1 (encoded by *MAP3K7*) to the inhibitor of NF-κB complex (IκB complex, encoded by *CHUK, IKBKB, IKBKG*), as strongly required for IFN-γ and to a lesser extent IL-2 production (Fig 2D-E and fig. S6D). These essential intracellular signaling components were not identified by the CRISPRa screens. In contrast, CRISPRa revealed a set of NF-κB signaling membrane proteins, including multiple TNFRSF receptors and IL1R1, which can be overexpressed to enhance IFN-γ production (Fig. 2D-E). These receptors bottleneck IFN-γ production, but are not individually required for the cytokine response and were missed by CRISPRi. CRISPRa and CRISPRi together reveal both the essential TCR - NF-κB signaling circuit and nonessential, but potent, TNFRSF receptors, as signaling components that regulate IFN-γ production.

We used our integrated dataset combined with literature review to build a high-resolution map of tunable regulators in TCR signal transduction to cytokine response (Fig. 2F). Strikingly, many of the screen hits could not be fit in with previous literature and are potential novel functional regulators of cytokine production (Fig. 2G). Many of the novel positive regulators of T cell cytokine response were only found in the CRISPRa screens (e.g. *APOBEC3A/D/C*, *FOXQ1,* and *EMP1*), underscoring the need for gain-of-function screens for comprehensive discovery. Thus, paired CRISPRa and CRISPRi screens complimentarily map the tunable genetic circuits controlling T cell stimulation responsive cytokine production.

Given the lack of previous unbiased gain-of-function studies in T cells and the robust identification of novel genes that may regulate T cell cytokine production, we next performed arrayed CRISPRa experiments for deeper phenotypic characterization of screen hits (Fig. 3A). We cloned a panel of 30 arrayed sgRNAs, targeting 14 screen hits, at two sgRNAs per gene (including one not present in the Calabrese library used for the screens) plus two non-targeting control sgRNAs. Target genes were selected from different screen categories, including IL-2 and IFN-γ shared negative, shared positive, or specific regulators of IL-2 or IFN-γ (Fig. 3B). We biased our selection to genes with little to no previous evidence of T cell or cytokine production function, however we included the established regulators *VAV1*, and *MAP4K1*, as well as the positive controls *IL2 and IFNG*. First, we validated that each of the sgRNAs caused robust increases in expression of target gene mRNA (fig. S7). Next, we assessed production of IL-2 and IFN-γ by intracellular staining. In this arrayed format, we also tested whether the regulators of IL-2 and IFN-γ identified in our CRISPRa screens may also affect another key cytokine, TNF-a. TNF-ɑ, like IL-2 and IFN-γ, is a Th1-associated cytokine, is secreted by CD4^+^ and CD8^+^ T cells, and has important roles in autoimmune disease and cancer (*29*). Flow cytometry demonstrated robust increases in all three cytokines in stimulated *VAV1* CRISPRa cells, compared to control cells (Fig. 3C). We completed intracellular cytokine staining on the full arrayed panel, using T cells from 4 screen-independent donors. Remarkably, we found that for 13 of 14 target genes, we were able to detect significant changes in the proportion of cells positive for the relevant cytokine(s), with at least one sgRNA and in most cases both, supporting the precision of the genome-wide screens (Fig. 3D and fig. S8). Of note, with the exception of *TNFRSF1A,* all positive regulators that increased stimulation-dependent cytokine production, did not cause spontaneous cytokine production in the absence of stimulation (Fig. 3D and fig. S8). Although IL-2 was screened in CD4^+^ cells, and IFN-γ in CD8^+^ cells, sgRNA effects were generally consistent across both T cell lineages. Overall, hits which were regulators of both IL-2 and IFN-γ, also tended to alter production of TNF-ɑ suggesting that these genes may act as general regulators of T cell response and may broadly alter cytokine response to stimulation (Fig. 3E).

**Figure 3.**
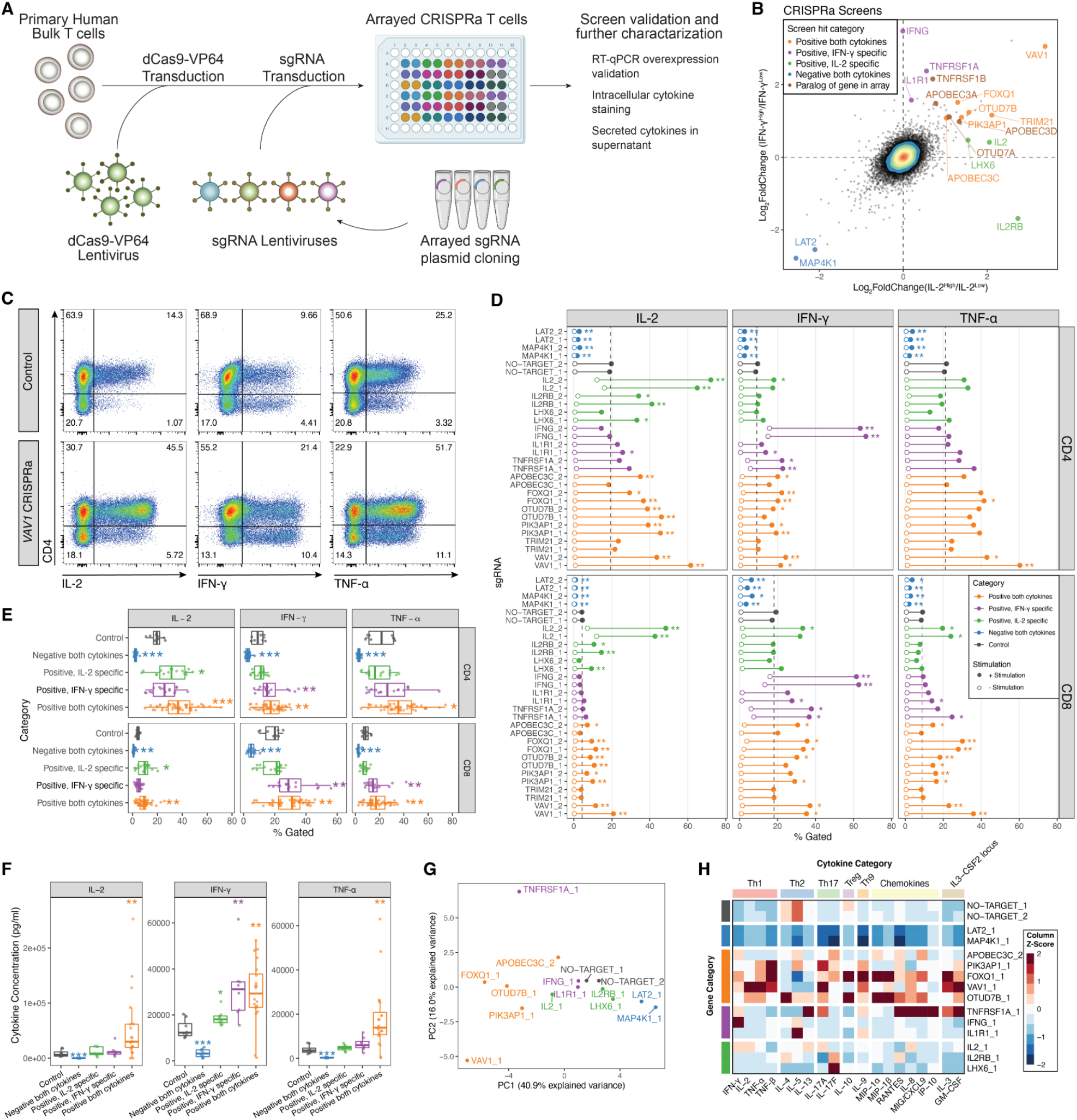
Further characterization of CRISPRa screen hits by arrayed profiling. **(A)** Schematic of arrayed experiments. **(B)** Comparison of IL-2 and IFN-γ CRISPRa screens, with genes targeted by the arrayed sgRNA panel indicated, as well as their screen hit categorization. Paralogs of arrayed panel genes that were also highly ranked hits are additionally indicated. **(C)** Intracellular cytokine staining flow cytometry for indicated cytokines in control (NO-TARGET_1 sgRNA) or *VAV1* (VAV1_1 sgRNA) CRISPRa T cells after 10 hours of stimulation. **(D)** Intracellular cytokine staining of full arrayed sgRNA panel, showing percent of cells gated positive for indicated cytokines in CD4^+^ or CD8^+^ T cells. Points represent the mean value of 4 donors, with and without stimulation. Dashed vertical lines represent the mean no-target control sgRNA control value with stimulation. *q<0.05, **q<0.01, Mann-Whitney U-test followed by q-value multiple comparison correction. Full data in fig. S8B. Medium stimulation dose shown for IL-2 and IFN-γ, low dose shown for TNF-ɑ. **(E)** Intracellular cytokine staining arrayed panel binned by indicated gene categories, with sgRNAs targeting *IL2* and *IFNG* removed. Points represent a single sgRNA and donor measurement. *p<0.05, **p<0.01, ***p<0.001, Mann-Whitney U-test. **(F)** Secreted cytokine staining arrayed panel grouped by indicated gene categories, with sgRNAs targeting *IL2* and *IFNG* genes removed. Points represent a single gene and donor measurement. *p<0.05, **p<0.01, ***p<0.001, Mann-Whitney U-test. **(G)** Principal component analysis of secreted cytokine measurements from indicated sgRNAs. **(H)** Heatmap of selected secreted cytokine measurements grouped by indicated biological category. Values represent the median of 4 donors, followed by z-score scaling for each cytokine.

We next tested if genes identified by CRISPRa could also regulate cytokines when overexpressed as cDNA transgenes, because continuous expression of CRISPRa would present challenges in cell therapies due to Cas9 immunogenicity (*30*) (fig. S9A). In an arrayed panel of lentivirally delivered transgenic cDNAs we observed gene overexpression effects on cytokine production generally consistent with CRISPRa (fig. S9B-C). The cDNA transgenes also functioned with more physiological antigen stimulation. Overexpression of these cDNA constructs in T cells transduced with a transgenic TCR that recognizes the NY-ESO-1 cancer antigen (1G4) led to similar effects on cytokine production when stimulated with NY-ESO-1^+^ cancer cells (fig. S9D). These findings suggest that overexpression of CRISPRa screen hits eventually could be used to reprogram functional responses in antigen-specific cell therapies.

We next assessed how individual CRISPRa perturbations reprogram cytokine responses by measuring a broad panel of secreted cytokines. Guided by the intracellular cytokine staining results, we selected one sgRNA per gene plus two non-targeting control sgRNAs, and performed multiplexed secretome measurements, consisting of 48 secreted cytokine and chemokines, 32 of which were detected in all 4 donors in control samples (fig. S10A). First, we examined IL-2, IFN-γ, and TNF-ɑ in the supernatants and found that sgRNA effects on secreted protein measurements were mostly consistent with intracellular staining (Fig. 3F and fig. S10B). Next, we analyzed the broad spectrum of cytokine measurements with principal component analysis (PCA) and hierarchical clustering, and observed sgRNA categorical grouping consistent with those observed in the screens, with sgRNAs targeting genes identified as regulators of both IL-2 and IFN-γ causing broad increases or decreases in cytokine concentration (Fig. 3G and fig. S10C). Interestingly, we discovered distinct patterns in the classes of cytokines increased by different regulators (Fig. 3H). sgRNAs targeting *VAV1* (a proximal positive regulator of TCR signaling) and *FOXQ1* (a poorly characterized transcription factor) led to preferential increases in Th1 signature cytokines and dampened Th2 cytokines. Surprisingly, activating *OTUD7B*, encoding another positive regulator of proximal TCR signaling (*31*), had a divergent effect from *VAV1* and led to a marked increase in Th2 cytokines and a lower magnitude increase in Th1 cytokines (fig. S10D). Taken together, this suggests that identified regulators may not only amplify TCR stimulation and signaling, but also tune the T cell secretome towards specific signatures.

CRISPRa should identify novel regulators of cytokine production, even if they are not normally expressed in T cells under the conditions tested. Indeed, while the CRISPRi hits were biased towards genes with high mRNA expression (as expected), CRISPRa hits followed a wider distribution including non-expressed genes (Fig. 4A). Included among the non-expressed or very low expressed genes were validated hits, including *IL1R1, PIK3AP1*, and genes encoding the transcription factors *FOXQ1* and *LHX6*, as shown by RNA-seq and RT-qPCR (Fig. 4A and fig. S7B). These genes could either be context-dependent regulators of T cell function that are not active under these cell culture conditions or regulators normally active in other cell types with new synthetic functions upon forced expression in T cells. The cytokine IL-1 has reported physiological functions in T cells *(32, 33)*, although its receptor, IL1R1, is not expressed at high mRNA levels in the T cell context tested here (Fig. 4A). CRISPRa of *IL1R1* may have sensitized the T cells to autocrine IL-1, and revealed its function as a regulator of IFN-γ. In contrast to *IL1R1*, which is expressed in T cells in specific contexts, *FOXQ1* is expressed primarily in non-immune epithelial cells and the transcript is barely detectable in most peripheral T cells (fig. S11) (*34*, *35*), but strikingly *FOXQ1* gain-of-function drove strong cytokine responses.

**Fig 4.**
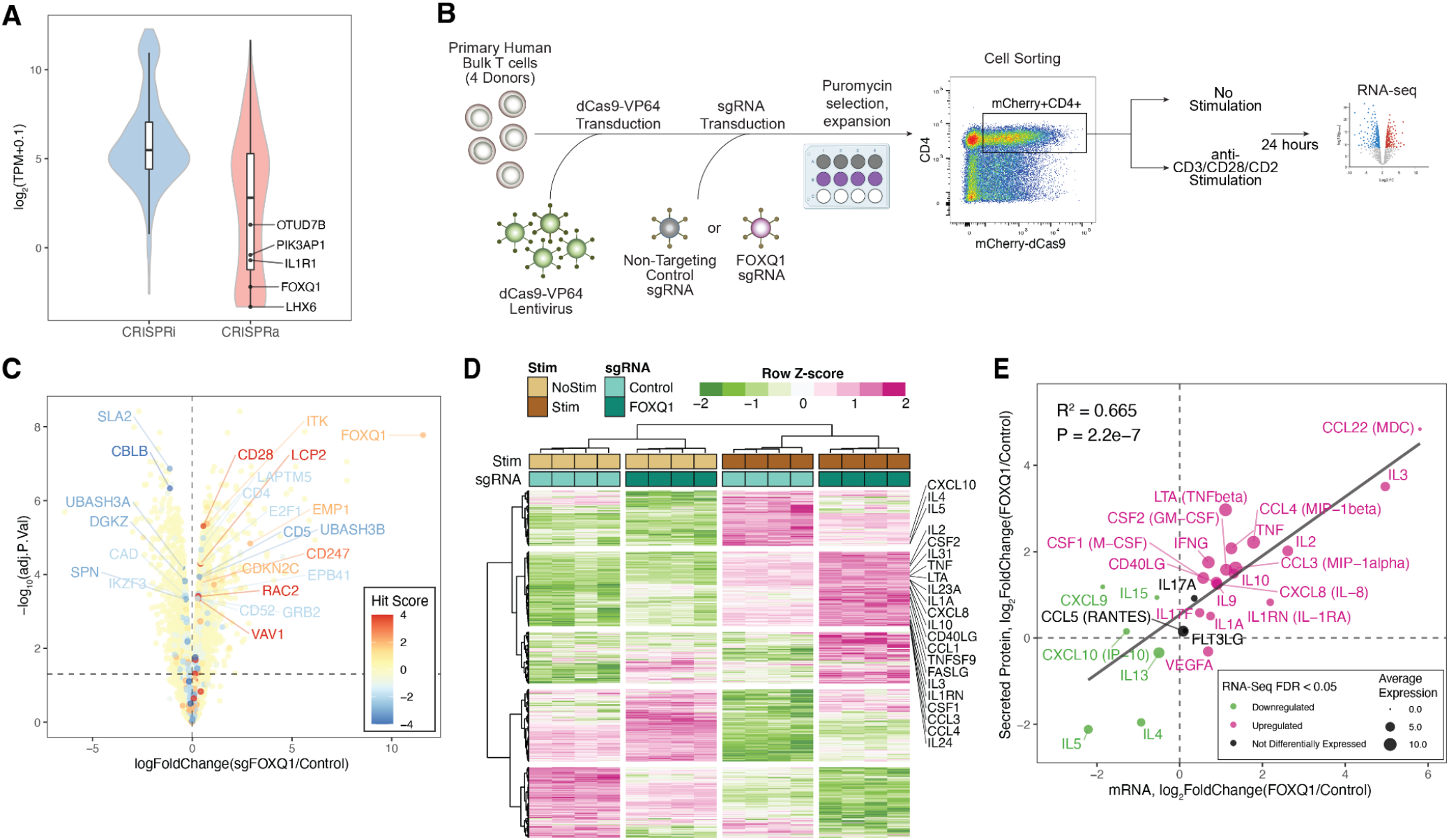
*FOXQ1* overexpression transcriptionally reshapes the cytokine response in CD4^+^ T cells. **(A)** Distributions of gene mRNA expression for CRISPRa and CRISPRi cytokine screen hits in resting CD4^+^ T cells. Genes from the arrayed CRISPRa panel with very low expression are indicated. **(B)** Schematic of RNA-seq experiment in CD4^+^ T cells after *FOXQ1* overexpression. **(C)** Differential expression volcano plot of *FOXQ1* sgRNA versus non-targeting control in non-stimulated cells. Points are colored according to hit score, where hit score is defined as the number of screens which identified the gene was identified as a positive regulator (positive hit score) or negative regulator (negative hit score) of cytokine production. **(D)** Heatmap representation of *FOXQ1* differentially expressed genes across all samples. Selected genes represent any that have an FDR adjusted p value < 1e-4 in *FOXQ1* sgRNA versus control differential expression tests. Selected cytokine genes of interest are indicated on the right. **(E)** Comparison of log_2_ fold changes in mRNA and secreted protein expression (Fig. 3H and fig. S10) caused by *FOXQ1* overexpression.

To investigate this “synthetic” function of FOXQ1 in human CD4^+^ T cells, we performed transcriptomic analysis with RNA-seq (Fig. 4B). Differential expression analysis revealed widespread gene expression changes in T cells overexpressing *FOXQ1* in both resting conditions (>3800 differentially expressed genes) and upon re-stimulation (>5500 differentially expressed genes) relative to control cells (fig. S12A-C). In resting T cells, *FOXQ1* overexpression increased expression of core cytokine regulators identified in the CRISPRa and CRISPRi screens (*CD28, LCP2, RAC2, VAV1*) and reduced expression of core negative regulators (*CBLB, SLA2*), suggesting FOXQ1 may sensitize T cells to stimulation by increasing overall levels of proximal signaling components (Fig. 4C). Unsupervised clustering revealed that FOXQ1 does not simply sensitize, but rather, remodels the T cell stimulation response. Interestingly, the expression of a cluster of stimulation-responsive genes, containing most cytokine encoding genes, including *IL2* and *TNF*, was increased by *FOXQ1* overexpression; whereas a separate cluster of stimulation-responsive genes containing genes encoding Th2 cytokines, *IL4* and *IL5*, was dampened (Fig. 4D and fig. S12D). This is consistent with *FOXQ1* causing a slight decrease in GATA3 expression and increase in *TBX21* (stimulation only), although it is unclear if the magnitude is enough to drive these differences (fig. S12E). Importantly, FOXQ1 driven differences in cytokine mRNA expression were consistent with secreted cytokine measurements, as shown by significant correlation (Fig. 4E). In conclusion, *FOXQ1* overexpression causes a reshaped stimulation response with increased transcription of most stimulation responsive cytokines.

Paired genome-wide CRISPRa and CRISPRi screens complement each other to decode the genetic programs regulating stimulation-responsive cytokine production in primary human T cells. CRISPRi enabled the identification of genes that are individually required (and not redundant) for cytokine production. CRISPRa identified key signaling bottlenecks in pathway function, and regulators that are not necessarily active in *ex-vivo* cultured T cells. These inactive regulators may represent genes expressed in rare, poorly characterized immune cell types, genes only active under certain immune cell circumstances, or genes canonically active in non-immune cell types but able to serve synthetic functions when experimentally induced in T cells.

Importantly, the technologies developed in this study enable new screening approaches in primary human T cells and potentially other primary cell types. CRISPRi and CRISPRa are powerful tools for efficient perturbation of noncoding elements at scale, with the potential to enable scalable noncoding screens in primary T cells (*17*, *36*, *37*). Furthermore, this screening framework should be adaptable to other non-cutting applications of the CRISPR toolkit (*38*), continuing to expand opportunities to interrogate complex biological questions in primary cells.

We expect this study – and future CRISPRa gain-of-function screens – to accelerate development of improved T cell therapies. Notably, recent preclinical work revealed c-JUN overexpression in CAR T cells as an approach to limit exhaustion (*39*), highlighting the promise for gene over-expression to enhance cell therapies. Although we do not expect all perturbations that lead to increased cytokine production to translate to enhanced *in vivo* antitumor efficacy, we are encouraged by the identification of known T cell therapy targets in these screens. In particular, the CRISPRa IFN-γ screen identified gene targets for cancer immunotherapies in various stages of development, such as *TNFRSF9* (encoding 4-1BB), *CD27, CD40, and TNFRSF4* (encoding OX40), suggesting high potential for further investigation of hits in preclinical models. Hits identified by these genome-wide gain-of-function studies highlight components - both positive and negative regulators - that could be incorporated into synthetic circuits for cell therapies. This work should help guide cell engineering to tune defined cytokine responses up or down for different clinical indications. Further screens for additional relevant T cell functions will continue to nominate targets that can be modulated in next-generation adoptive cellular therapies.

## Acknowledgements

We thank the Parnassus Flow Cytometry Core Facility (PFCC) for support with sorting. We thank Eric Shifrut, Julia Carnevale, and Victoria Tobin for help related to the development of CRISPRa technologies in T cells. We thank Justin Eyquem for providing NY-ESO-1 expressing NALM6 cells. Zachary Steinhart is supported by the Parker Institute for Cancer Immunology scholarship. Ralf Schmidt was supported by fellowships of the Austrian Exchange Service and the Austrian Society of Laboratory Medicine. We thank all members of the Marson lab for critical insight and discussion over the course of this study.

## Funding

National Institute of Diabetes and Digestive and Kidney Diseases DP3DK111914-01 (AM)

Simons Foundation (AM)

Burroughs Wellcome Fund, Career Award for Medical Scientists (AM)

The Cancer Research Institute (CRI)

Lloyd J. Old STAR grant (AM)

Parker Institute for Cancer Immunotherapy, PICI (AM)

Innovative Genomics Institute (IGI)

National Institutes of Health grant P30 DK063720 (Parnassus Flow Cytometry Core, AM)

National Institutes of Health grant S10 1S10OD021822-01 (Parnassus Flow Cytometry Core, AM)

Investigator at the Chan Zuckerberg Biohub (AM)

Gifts from Brook Byers, Barbara Bakar, Karen Jordan, Elena Radutzky

## Author contributions

Conceptualization: RS, ZS, AM

Methodology: RS, ZS, ML, FB, VQN

Investigation: RS, ZS, JWF, AM

Visualization: RS, ZS

Funding acquisition: RS, ZS, AM

Project administration: RS, ZS, AM

Supervision: AM

Writing – original draft: RS, ZS, AM

Writing – review & editing: RS, ZS, ML, JWF, VQN, FB, AM

## Competing interests

A.M. is a compensated co-founder, member of the boards of directors, and a member of the scientific advisory boards of Spotlight Therapeutics and Arsenal Biosciences. A.M. was a compensated member of the scientific advisory board at PACT Pharma and was a compensated advisor to Juno Therapeutics and Trizell. A.M. owns stock in Arsenal Biosciences, Spotlight Therapeutics, and PACT Pharma. A.M. has received honoraria from Merck and Vertex, a consulting fee from AlphaSights, and is an investor in and informal advisor to Offline Ventures. The Marson lab has received research support from Juno Therapeutics, Epinomics, Sanofi, GlaxoSmithKline, Gilead, and Anthem. R.S., Z.S., and A. M. are listed as inventors on a patent application related to this work.

## Data and materials availability

All raw sequencing data critical to the findings of this study, including CRISPR screens and RNA-seq, are deposited at NCBI Gene Expression Omnibus under accession number GSE174292.

## Materials and Methods

### Isolation and culture of human T cells

Human T cells were sourced from PBMC enriched leukapheresis products (Leukopaks) from healthy donors, following informed written consent (Stemcell Technologies). Bulk T cells were isolated from Leukopaks using EasySep magnetic selection following manufacturers’ recommended protocol (Stemcell Technologies cat 17951). Immediately following isolation, bulk T cells were frozen in Bambanker Cell Freezing Medium at 50e6 cells/ml (Bulldog Bio cat BB01) and stored at −80°C for short term or liquid nitrogen for long term storage. Unless otherwise noted, thawed T cells were cultured in X-VIVO 15 (Lonza Bioscience cat 04-418Q) supplemented with 5% FCS, 55mM 2-mercaptoethanol, 4mM N-Acetyl L-Cysteine, and 500 IU/ml recombinant human IL-2 (Amerisource Bergen cat 10101641). Primary T cell activation was done by anti-humanCD3/CD28 CTS dynabeads (Fisher Scientific cat 40203D) at a 1:1 cell to bead ratio at 1e6 cells/ml.

### Cell line maintenance

Lenti-X HEK293T cells (Takara Bio cat 632180) were maintained in DMEM high glucose with GlutaMAX™ (Fisher Scientific cat 10566024), supplemented with 10% FCS, 100U/ml PenStrep (Fisher Scientific cat 15140122), 1mM Sodium Pyruvate (Fisher Scientific cat 11360070), 1x MEM Non-Essential Amino Acids (Fisher Scientific cat 11140050) and 10mM HEPES solution (Sigma cat H0887-100ML). Cells were passaged every two days using Tryple Express (Fisher Scientific cat 12604013) for dissociation and kept at a confluency of <60%.

NALM6 cells were engineered to express NY-ESO-1 peptide in an HLA-A0201 background, recognizable with the 1G4 TCR by the Eyquem lab at UCSF and provided for TCR stimulation co-culture experiments. For simplicity, these cells are referred to as NALM6. NALM6 cells were cultured in RPMI (Gibco cat 21870092) supplemented with 10% FCS, 100U/ml PenStrep (Fisher Scientific cat 15140122), 1mM Sodium Pyruvate (Fisher Scientific cat 11360070) and 1x MEM Non-Essential Amino Acids (Fisher Scientific cat 11140050), 10mM HEPES solution (Sigma cat H0887-100ML) and 2mM L-Glutamine (Lonza Bioscience cat 17-605E).

### Plasmids

dCas9-VP64 originated from lentiSAMv2 (addgene 75112) and cloned into the lentiCRISPRv2-dCas9 backbone (addgene 112233) with Gibson Assembly. The promoter was switched to SFFV and mCherry was introduced upstream of dCas9-VP64, separated by a P2A sequence resulting in the pZR112 plasmid. The LTR-LTR range was minimized to enhance lentiviral titer. For CRISPRi, BFP in pHR-SFFV-dCas9-BFP-KRAB (addgene 46911) was switched to mCherry with Gibson Assembly resulting in pZR244.

Single sgRNAs for arrayed experiments have been introduced by Golden Gate Cloning as described before(*20*). Briefly, DNA oligomers with Golden Gate overhangs were annealed and subsequently cloned into the non-digested target plasmid using the NEB^®^ Golden Gate Assembly Kit (BsmBI-v2, New England Biolabs cat E1602L). sgRNAs have been cloned into pXPR_502 (addgene 96923) for CRISPRa and into CROPseq-Guide-Puro(*40*) (addgene 86708) for CRISPRi. All single sgRNAs used in this study are found in table S3.

The genome wide CRISPRa (Calabrese A 92379 and Calabrese B 92380) and CRISPRi libraries (Dolcetto A 92385 and Dolcetto B 92386) were obtained from addgene. 40 nanograms of each library were transformed into Endura™ ElectroCompetent Cells (Lucigen cat 60242-2) following the manufacturer’s instructions. After transformation, Endura cells were grown in a shaking incubator for 16h at 30°C in the presence of Ampicillin. Library plasmid has been isolated using the Qiagen Plasmid Plus MaxiKit (Qiagen 12963) and sequenced for sgRNA representation as described under “Genome-wide CRISPRa and CRISPRi screens”.

For cDNA mediated target overexpression, the lentiCRISPRv2 (addgene 75112) backbone was rebuilt to a lentiviral cDNA cloning plasmid with an SFFV promoter followed by BsmBI restriction sites and P2A-Puro. Transgene cDNAs were purchased from Genscript, choosing the canonical (longest) isoform for each gene, and BsmBI restriction sites were introduced by PCR. The final lentiviral transfer plasmids were assembled using the NEB^®^ Golden Gate Assembly Kit (BsmBI-v2, New England Biolabs cat E1602L).

### Lentivirus production

Unless otherwise stated, HEK293T cells were seeded in Opti-MEM™ I Reduced Serum Medium (OPTI-MEM) with GlutaMAX™ Supplement (Gibco cat 31985088) supplemented with 5% FCS, 1mM Sodium Pyruvate (Fisher Scientific) and 1x MEM Non-Essential Amino Acids (Fisher Scientific) (cOPTI-MEM) at 36e6 cells per T225 flask in 45ml of medium overnight to achieve confluency between 85% and 95% at the time point of transfection. Next morning, HEK293Ts cells were transfected with 2nd generation lentiviral packaging plasmids and transfer plasmid using Lipofectamine 3000 transfection reagent (Fisher Scientific cat L3000075). In detail, 165ul of Lipofectamine 3000 reagent was added to 5ml room-temperature OPTI-MEM without supplements. 42μg of Cas9 transfer plasmid, 30μg psPAX2 (addgene 12260), 13μg pMD2.G (addgene 12259), and 145μl p3000 reagent were added to 5ml room temperature OPTI-MEM without supplements and mixed by gentle inversion. The plasmid and Lipofectamine 3000 mixes were combined, mixed by gentle inversion, and incubated for 15 minutes at room temperature. Following incubation, 20ml of medium were removed from the T225 flask and the 10ml transfection mixture was carefully added without detaching HEK293T cells. After 6 hours, the transfection medium was replaced with 45ml cOPTI-MEM supplemented with 1x ViralBoost (Alstem Bio cat VB100). Lentiviral supernatant was harvested 24hr after transfection (first harvest) and replaced with 45ml fresh cOPTI-MEM. A second harvest was done 48hr after transfection. Immediately after collection, the media was spun down at 500xG, 5 minutes, 4°C, to clear cellular debris. Unless otherwise noted, Lenti-X-Concentrator (Takara Bio 631232) was added to the collected supernatant and lentivirus was concentrated following the manufacturer’s instructions and resuspended in OPTI-MEM in 1/100th volume of the original culture volume without supplements. Lentiviral particles were subsequently aliquoted and frozen at −80°C.

### Genome-wide CRISPRa and CRISPRi screens

One day after activation, T cells from 2 Donors were infected with 2% v/v concentrated dCas9-VP64 lentivirus. Two days after activation, T cells were split into two populations and infected with 1% v/v (MOI ~0.5) Calabrese Set A (addgene 92379) or 0.8% v/v (MOI ~0.5) Calabrese Set B (addgene 92380) lentivirus. These two sets were independently cultured and processed in parallel until analysis. Three days following activation, fresh media with IL-2 (final concentration 500IU/ml) and puromycin (final concentration 2μg/ml) was added to bring cells to 0.3e6 cells/ml. Cells were split two days later and fresh media with IL-2 was added to bring cells to 0.3e6 cells/ml. Two days later, fresh media without IL-2 was added to bring cells to 1e6/ml. Eight days after initial activation, cells were harvested, spun down and resuspended at 2e6 cells/ml X-VIVO 15 without supplements. Next day, cells were restimulated and stained for FACS as described under intracellular cytokine staining. The following two days, cells were sorted at the Parnassus Flow Cytometry Core Facility (PFCC) in IL-2 low and high CD4^+^ T cells and IFN-γ low and high CD4^−^ T cells (see Figure S3C for gating strategy). Sorted cells were stored in EasySep Buffer (PBS with 2% FCS and 1mM EDTA) overnight until genomic DNA isolation.

The same experimental procedure using T cells from the same donors was followed for the CRISPRi screens. T cells were infected with dCas9-mCherry-KRAB at 2% v/v and Dolcetto A (addgene 92385) and B (addgene 92386) sgRNA libraries at 10 % v/v or 25% v/v unconcentrated virus, respectively (~0.5 MOI).

Genomic DNA was extracted from fixed cells as described previously (*41*). Integrated sgRNA sequences were amplified as previously described (*20*), and sequencing libraries were subsequently agarose gel purified using NucleoSpin Gel and PCR Clean‑up Mini kit (Machery-Nagel cat 740609.50). Libraries were sequenced on a NextSeq500 instrument to a targeted depth of 100-fold coverage.

### CRISPR screen analysis

Reads were aligned to the appropriate reference library using MAGeCK version 0.5.9.2 (*42*) using --trim-5 22,23,24,25,26,28,29,30 argument to remove the staggered 5’ adapter. Next, raw read counts across both library sets were normalized to the total read count in each sample and each of the matching samples across two sets were merged in order to generate a single normalized read count table. Normalized read counts in high versus low bins were compared using *mageck test* with *--norm-method none*, *--paired,* and *--control-sgrna* options, pairing samples by donor and using non-targeting sgRNAs as controls, respectively. Gene hits were classified as having a median absolute log2FoldChange value greater than 0.5 and an FDR < 0.05.

### Gene set enrichment analysis

Gene set enrichment analysis was completed with the fgsea Bioconductor R package using default settings (*43*). KEGG pathways v7.4 were obtained from GSEA mSigDB http://www.gsea-msigdb.org/gsea/downloads.jsp. The KEGG NF-kappa B signaling pathway (entry hsa04064) was missing from this dataset and added manually from https://www.genome.jp/entry/pathway+hsa04064.

### Arrayed CRISPRa experiments

For each gene chosen to target in follow up experiments, one sgRNA was chosen from the Calabrese library used in screens. The first sgRNAs (“_1”) were manually chosen for consistent log2 fold change observed in both donors. The second sgRNA (“_2”) was picked from the hCRISPRa-v2 genome-wide library (*44*), choosing the top ranked sgRNA not present in Calabrese libraries for each gene. sgRNAs were cloned into the pXPR_502 vector as described in the plasmid section.

Primary human T cells were transduced with 2% v/v mCherry-2A-dCas9-VP64 lentivirus (pZR112) one-day post-activation. The following day (day 2), the dCas9-VP64 transduced cells were split into 96-well flat-bottom plates, avoiding edge wells, and transduced with a different sgRNA lentivirus in each well (5% v/v). One day after sgRNA transduction, fresh medium was added with IL-2 (500U/ml) and 2μg/ml puromycin (final culture concentrations). Cells were passaged two days later, adding fresh medium with 500U/ml IL-2 and maintaining a concentration of 0.3e6-1e6 cells/ml, and 96-well plates were copied as needed to maintain this concentration. On day 8 cells from copied plates were pooled and samples were counted. Cells were pelleted and resuspended at a concentration of 2e6 cells/ml in fresh X-VIVO-15 without additives. On day 9 cells were restimulated with anti-CD3/CD28/CD2 immunocult or left unstimulated.

### cDNA experiments

See Fig. S9A for experimental overview. One day after activation, T cells were transduced with the 1G4 TCR lentivirus recognizing the NY-ESO-1 antigen or non-transduced for immunocult assay. One day later, cells were transduced with the transgenes in cDNA format. Three days after initial activation, puromycin was added to obtain a final concentration of 2ug/ml along with fresh X-VIVO 15 media with 500IU/ml IL-2 and further cultured and expanded analogous to the genome wide CRISPR screens. Nine days after initial activation, T cells were spun down and resuspended at 2e6 cells/ml in X-Vivo 15 without supplements. On the same day, 1G4 TCR expression was assessed by flow cytometry following dextramer staining (Immudex cat WB3247-PE) to ensure even expression across different cDNA constructs. Next day, T cells were restimulated with either 6.25ul /ml Immunocult or NALM6 cells at an Effector: Target ratio of 1:2 for 1G4 TCR transduced cells. Cells were further processed as described under Intracellular cytokine staining. CD22 was used as a marker for NALM6 cells to discriminate from T cells in the co-culture. Overexpression of *OTUD7B* cDNA in together with the 1G4 TCR but not alone caused toxicity and was therefore excluded from analyses.

### Intracellular cytokine staining

Unless indicated otherwise, T cells were stimulated with ImmunoCult™ Human CD3/CD28/CD2 T Cell Activator (Stemcell Technologies cat 10990) with 6.25μl per ml culture media at 2e6cells/ml. One hour after re-stimulation, Golgi Plug protein transport inhibitor (BD Biosciences, cat 555029) was added at a 1/1000 dilution. Nine hours after addition of Golgi Plug, T cells were stained for surface antigens prior to fixation and subsequently processed for intracellular cytokine staining following BD Cytofix/Cytoperm™ kit (BD Biosciences cat 554714) instructions.

### Flow cytometry

Aria 2, Aria 3 and Aria Fusion cell sorters at the UCSF Parnassus Flow Core and the Gladstone Institute Flow Core were used (BD Biosciences) for sorting. The Attune NxT flow cytometer was used for flow cytometry. All antibodies used for flow cytometric analyses and sorting are summarized in table S4.

### Cytokine Luminex assay

T cells were prepared as explained under Arrayed CRISPRa experiments. On day 9 after activation, T cells at a concentration of 2e6/ml were re-stimulated with ImmunoCult™ Human CD3/CD28/CD2 (Stemcell Technologies cat 10970) at 6.25μl/ml. Twenty-four hours after re-stimulation, supernatant was collected and frozen at −20°C. Following a serial pilot titration, cytokine analyses were performed at a 1/200 dilution by Eve Technologies with the Luminex xMAP technology on the Luminex 200 system (Luminex). To remove very low expressed cytokines for downstream analysis, any group where 3 of 4 donors had undetectable cytokines, the cytokine was removed. Additionally, the sgIL1R1-1 Donor 4 measurement for IL-1alpha was manually removed, as this was an extremely high outlier.

### RT-qPCR

T cells were prepared as described under Arrayed CRISPRa experiments. Seven days post sgRNA transduction 100,000 T cells per well were pelleted at 500xG, 5 minutes, 4°C. Cells were lysed and RNA was extracted using Quick-RNA 96 kit (Zymo Research), following manufacturer’s protocol, skipping the option in-well DNase treatment. DNase treatment and cDNA synthesis were subsequently completed with Maxima First Strand cDNA Synthesis Kit for RT-qPCR, with dsDNase (Thermofisher Scientific). qPCR was performed with PrimeTime PCR Master Mix (Integrated DNA technologies) and PrimeTime qPCR probe assays (Integrated DNA Technologies, list of probes used in table S5) on a Applied Biosystems Quantstudio 5 real-time PCR system. Data was analyzed using the deltaDeltaCt method. The mean Ct values of two housekeeping genes, *PPIA* and *GUSB,* to calculate the deltaCt, and the mean deltaCt of non-targeting controls to calculate deltaDeltaCt.

### RNA-seq sample preparation

FOXQ1 and non-targeting sgRNA control primary human T cells from 4 donors were transduced and expanded as described in “Arrayed CRISPRa experiments” section. On day 8 mCherry+CD4+ populations were sorted and resuspended in X-VIVO-15 without additives at 2e6 cells/ml. On day 9 cells were restimulated with 6.25ul/ml anti-CD3/CD28/CD2 immunocult or left unperturbed for unstimulated condition. 24 hours post-stimulation cells were lysed for RNA-seq.

For RNA-seq, cells were lysed and RNA was purified using Quick-RNA Microprep kit (Zymo Research) without the optional in-well DNase treatment step. Purified RNA was treated with TURBO DNase (Thermofisher Scientific) to remove potential contaminating DNA. RNA was subsequently purified using RNA Clean & Concentrator-5 kit (Zymo Research). RNA quality control was performed using an RNA ScreenTape assay (Agilent), with all samples having a RNA integrity number > 7. RNA-seq libraries were prepared using the Illumina Stranded mRNA Prep kit, with 100ng of input RNA. Libraries were sequenced using paired end 72bp reads on a NextSeq500 instrument to an average depth of 32e6 clusters per sample.

### RNA-seq data analysis

Adapters were trimmed from fastq files using cutadapt version 2.10 (*45*) with default settings keeping a minimum read length of 20 bp. Reads were mapped to the human genome GRCh38 keeping only uniquely mapping reads using STAR version 2.7.5b (*46*) with the following settings “--outFilterMultimapNmax 1”. Reads overlapping genes were then counted using featureCounts version 2.0.1 (*47*) with the following settings “-s 2” and using the Gencode version 35 basic transcriptome annotation.

The count matrix was imported into R. Only genes with at least 1 count per million (CPM) across at least 4 samples were kept. TMM normalized counts were used for heatmaps. Differentially expressed genes between FOXQ1 overexpression and control samples were then identified using limma version 3.44.3 (*48*) while controlling for any differences between donors. Significant differentially expressed genes were defined as having an FDR adjusted p-value < 0.05.

### Statistical analysis

All statistical analyses were performed in R version 4.0.2, unless otherwise noted. In order to deal with ties in non-parametric tests, Mann-Whitney U-tests were performed using the wilcox_test function of the Coin R package (version 1.4-1), with default arguments. For q-value based multiple comparison correction, the R qvalue package (version 2.20.0) was used with default arguments.

## Supplemental Figures

**Figure S1.**
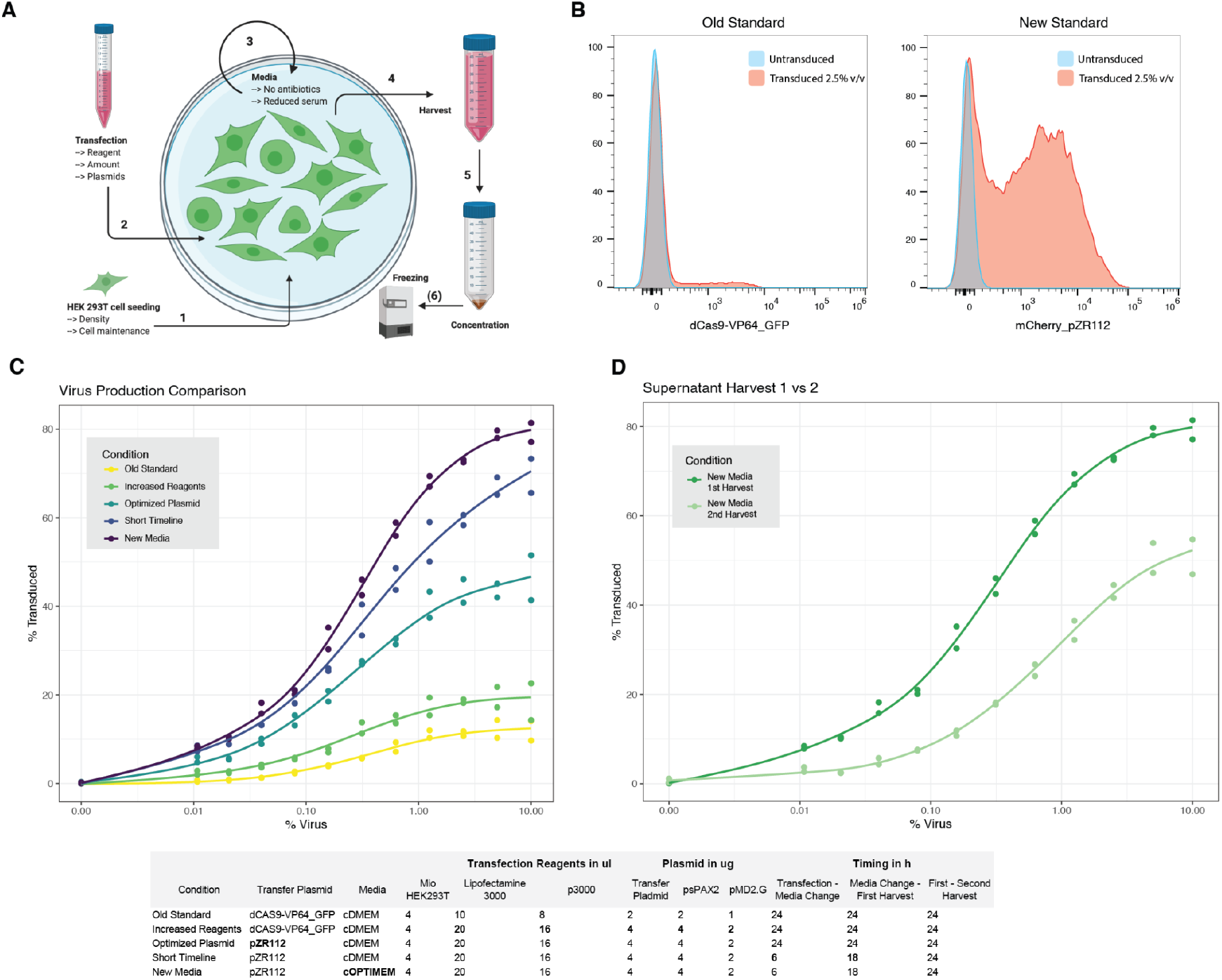
Optimization of lentiviral production and transduction in primary human T cells. **(A)** Schematic of factors leading to increased lentiviral titers in production protocols, Created with BioRender.com. **(B)** Representative example of transduction using previous vectors and protocols, versus after optimization as measured by flow cytometry. **(C)** Dose-response characterization of transduction efficiencies in primary T cells with indicated factors in lentiviral production. The table below contains full details for each condition. Each point represents % gated positive for fluorescent marker. N = 2 donors. **(D)** Comparison of transductions after harvesting lentivirus at indicated time points. N = 2 donors.

**Figure S2.**
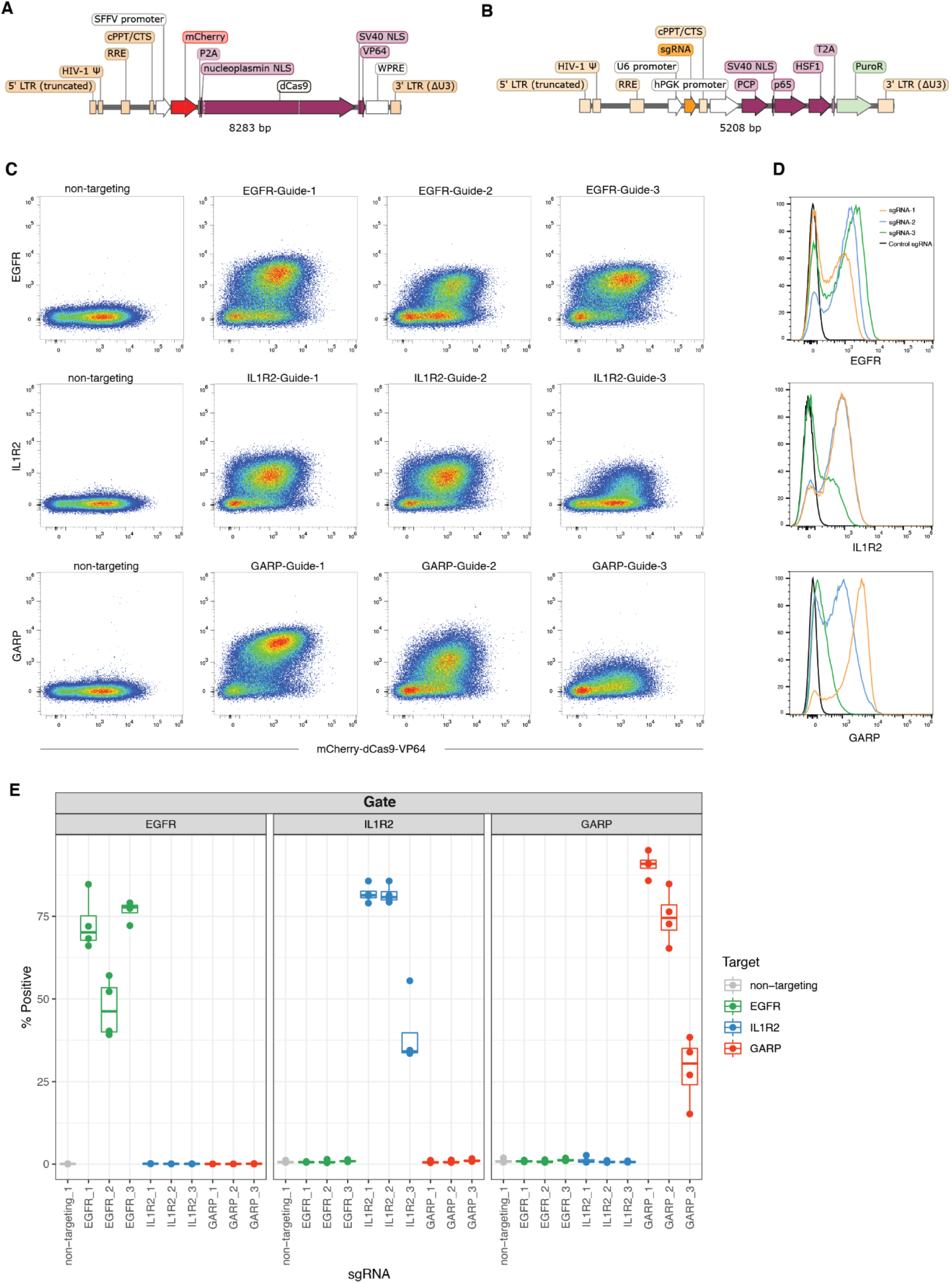
CRISPRa-SAM system enables robust gene activation in primary human T cells. **(A-B)** Lentiviral constructs used for dCas9-VP64 (A) and sgRNA, PCP-P65-HSF1 delivery (B). **(C-D)** Flow cytometry surface staining after delivery of sgRNAs targeting indicated genes or a non-targeting control sgRNA. Scatter plots showing surface marker correlation with mCherry-2A-dCas9-VP64 expression are shown in (C) and histograms in (D). **(E)** Quantification of (D) by gating. Points represent percent of cells gated in each donor (N = 4 donors).

**Figure S3.**
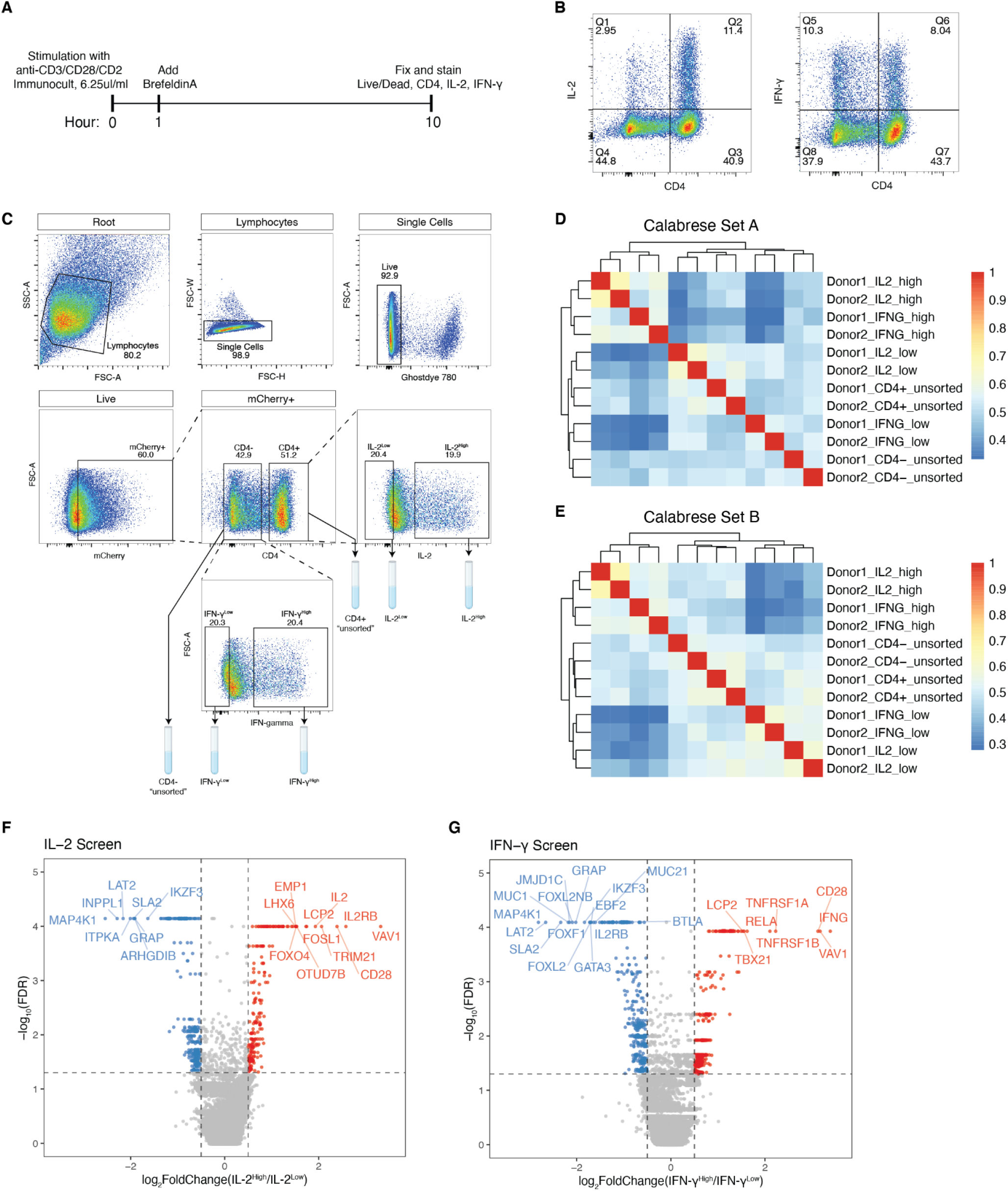
Genome-wide CRISPRa screens for cytokine production in primary human T cells. **(A)** Timeline of stimulation, golgi plug with Brefeldin A, and fixation/staining. **(B)** Representative flow cytometry plots showing CD4 staining compared to IL-2 and IFN-γ staining. **(C)** Gating strategy used for sorting screens with representative FACS plots. **(D-E)** Pairwise Pearson correlation matrices for Calabrese Set A (D) and Calabrese Set B (E) library samples. Correlations are calculated using sgRNA log_2_ fold change from the original plasmid pool. **(F-G)** Volcano plots for IL-2 (F) and IFN-γ (G) screens. Points represent each gene’s median log_2_ fold change followed by averaging two donors. Dashed lines represent cut-offs for hit calling, with positive and negative hits colored in red or blue, respectively.

**Figure S4.**
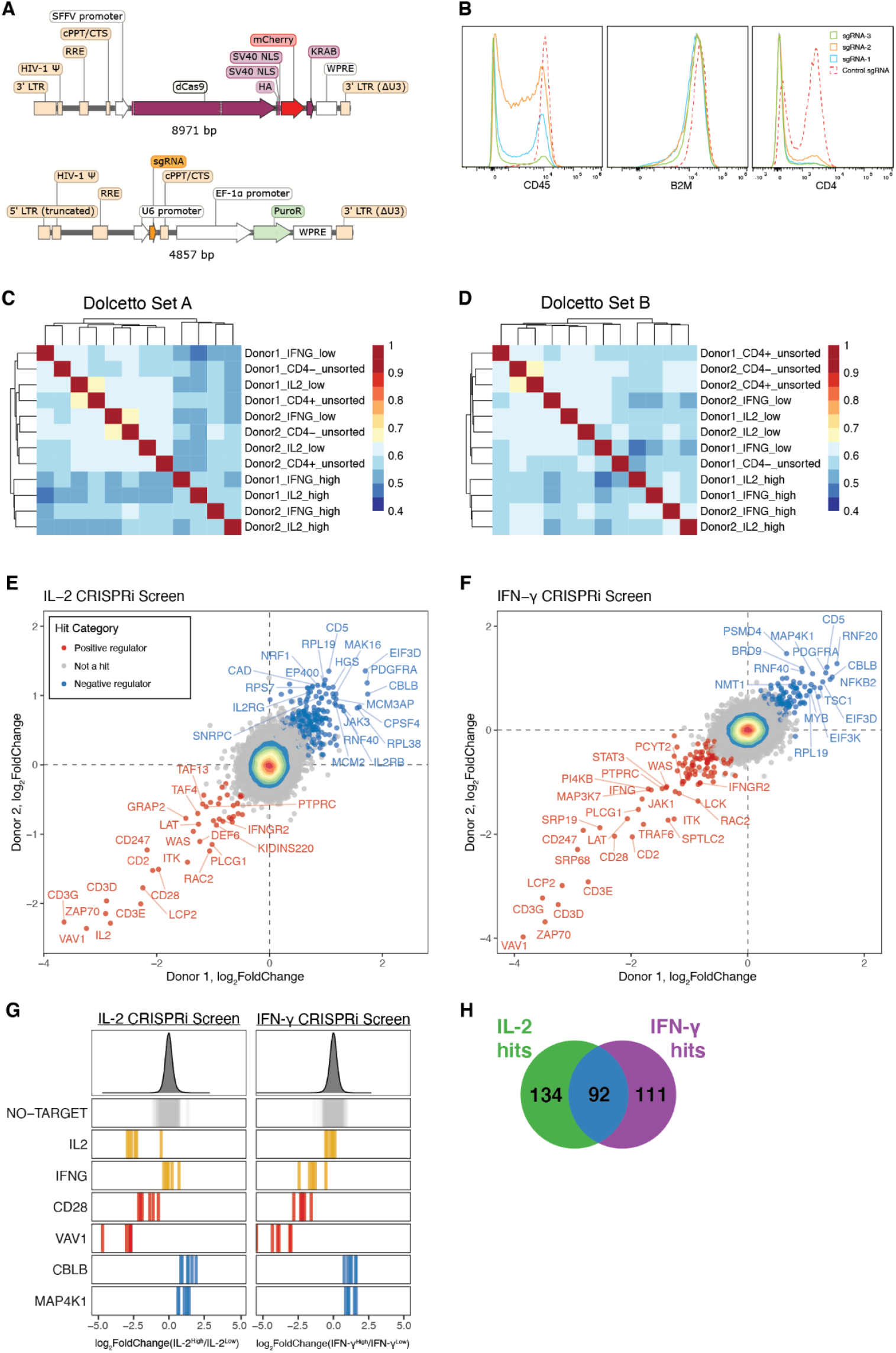
Genome-wide CRISPRi screens for cytokine production in primary human T cells. **(A)** Lentiviral constructs used for dCas9-KRAB and sgRNA delivery. **(B)** Representative histograms of indicated cell surface protein knockdown after delivery of sgRNAs targeting their transcriptional start site or a non-targeting control. **(C-D)** Pairwise Pearson correlation matrices for Dolcetto Set A (C) and Dolcetto Set B (D) library samples. Correlations are calculated using sgRNA log_2_ fold change from the original plasmid pool. **(E-F)** Scatter plots of median sgRNA log2 fold change (high/low sorting bins) for each gene, comparing screens in two donors, for IL-2 (E) and IFN-γ screens (F). Positive regulator hits are colored in red and negative regulator hits in blue. **(G)** sgRNA log_2_ fold changes for genes of interest in IL-2 (left) and IFN-gamma (right) screens. Bars represent the mean log_2_ fold change for each sgRNA across two donors. Density plots above represent the distribution of all sgRNAs. **(H)** Venn-diagram shows the number of screen unique and shared hits.

**Figure S5.**
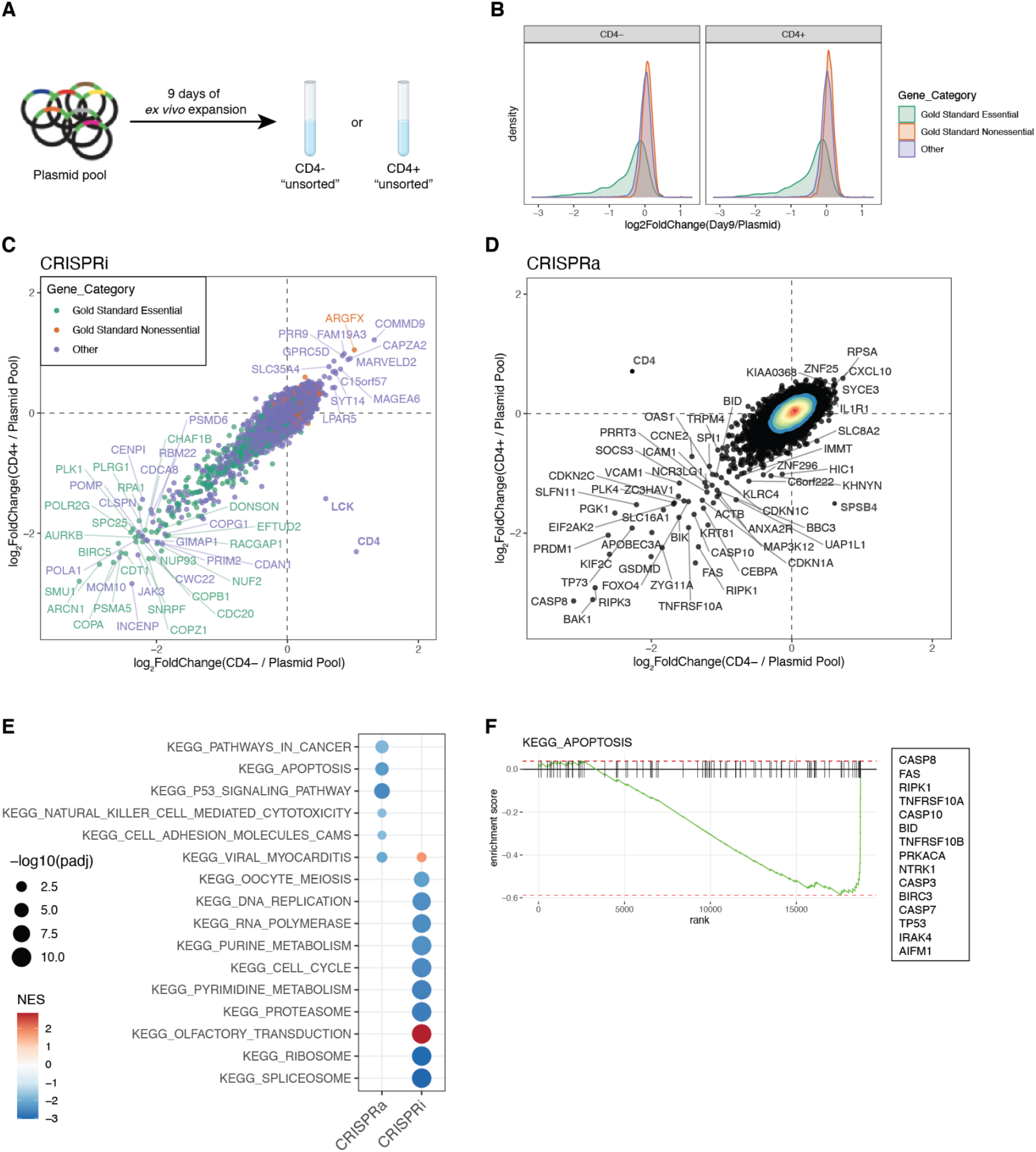
CRISPRi and CRISPRa dropout screens identify fitness genes. **(A)** Schematic of samples used for dropout analysis. **(B)** Log_2_ fold change distributions of sgRNAs targeting gold-standard essential and nonessential genes in CRISPRi screens. **(C-D)** Gene scatter plot of log_2_ fold change from plasmid pool to CD4^+^ and CD4^−^ “unsorted” populations in CRIPSRi and CRISPRa (D) screens. Points represent median of sgRNAs and mean of two donors. Notably, the vast majority of genes are highly correlated, with just *CD4, LCK*, and *SPSB4* discordant (bolded). The *CD4* difference is an artifact of the sorting strategy. Perturbations affecting *LCK*, and *SPSB4* may cause loss of CD4 expression. **(E)** GSEA of KEGG pathways significantly enriched in CRISPRa and CRISPRi dropout screens. Points are scaled to −log_10_ FDR adjusted P-value. **(F)** GSEA of KEGG Apoptosis pathway in CRISPRa dropout. Top ranked genes in the set are listed on the right.

**Figure S6.**
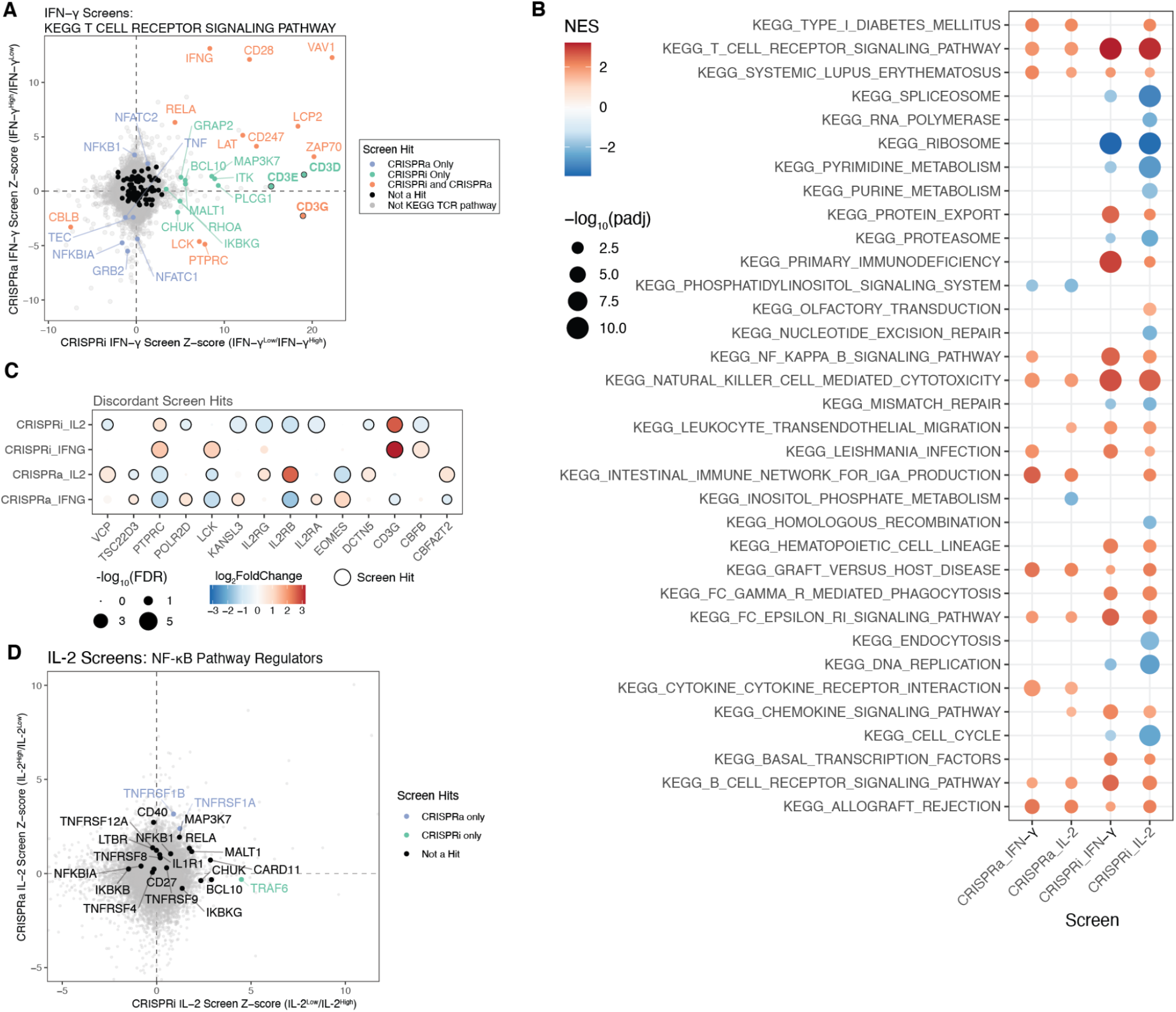
Pathway gene set enrichment in CRISPRa and CRISPRi cytokine screens. **(A)** Comparison IFN-γ CRISPRi and CRISPRa screens with genes belonging to the T cell receptor signaling pathway (KEGG pathways) indicated in colors other than grey. Genes meeting hit criteria are labeled. CD3 surface complex genes are indicated by black border and bolded labels. Scales represent z-score of log_2_ fold change, with positive regulators of IFN-γ production having positive z-scores. **(B)** Significantly enriched KEGG pathways from CRISPRa/CRISPRi screen log_2_ fold change gene ranks. Points are scaled according to −log_10_ FDR adjusted P-value. **(C)** Discordant hits across CRISPRi and CRISPRa hits, where perturbations identified as positive regulators are colored and red, and negative regulators in blue. Discordant hits between IL-2 and IFN-γ screens include *EOMES* and *CBFB*, encoding transcription factors known to have key roles in the differentiation of T cell subsets. The proximal kinase responsible for CD3ζ phosphorylation, LCK, and its activating phosphatase, CD45 (encoded by *PTPRC*), are discordant between CRISPRi and CRISPRa screens, suggesting appropriately balanced expression in this module is critical for optimal TCR signal transduction. **(D)** Comparison IL-2 CRISPRi and CRISPRa screens with genes of interest labeled.

**Supplemental Figure 7.**
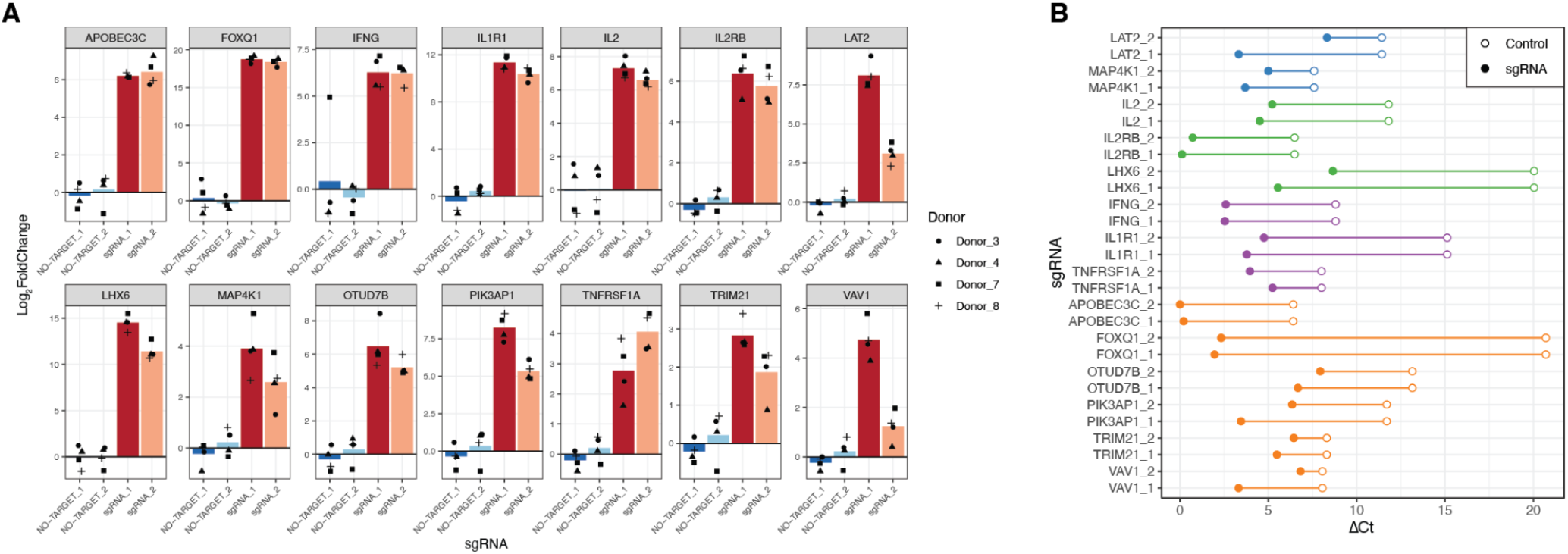
RT-qPCR validation of on-target gene activation in an arrayed CRISPRa panel. **(A)** Log_2_ fold change in mRNA expression for target sgRNAs compared to mean of non-targeting controls. Each facet represents the measurement of the indicated transcript and its measurement with two non-targeting control sgRNAs or two sgRNAs targeting its TSS. **(B)** deltaCt summary of (A), showing mean non-targeting control and mean targeting sgRNA deltaCt values for each gene.

**Figure S8.**
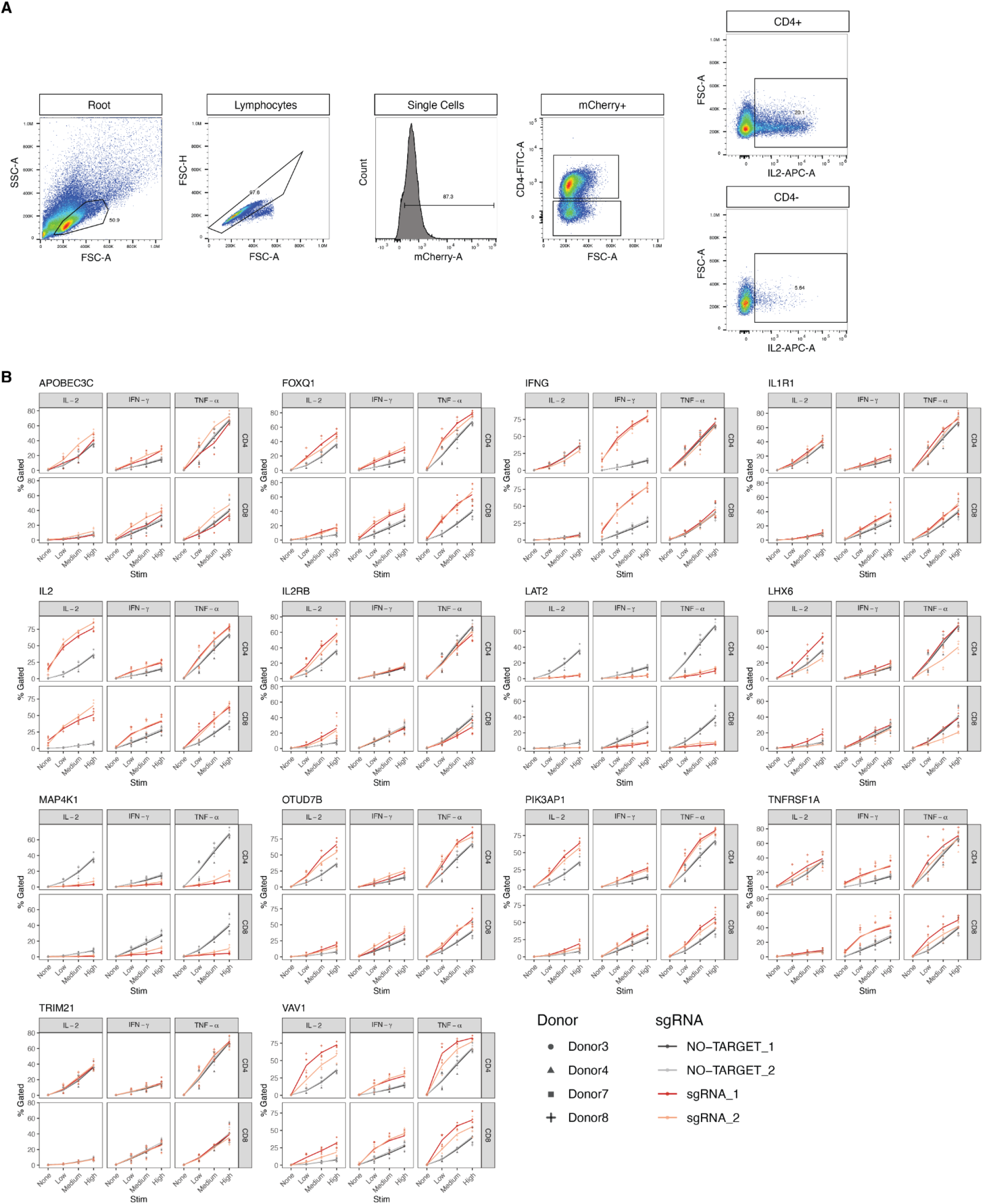
Intracellular cytokine staining in an arrayed CRISPRa panel. **(A)** Schematic of gating strategy. Note, IFN-γ and TNF-ɑ are not shown, but follow the same strategy as IL-2. **(B)** Complete data of summary shown in Fig. 3D. Points represent the percentage of cells gated as positive for expressing a given cytokine in the indicated donors, across multiple stimulation doses. Low, medium, and high doses represent 3.125, 6.25, and 12.5μl/ml of anti-CD2/CD28/CD2 Immunocult, respectively.

**Figure S9.**
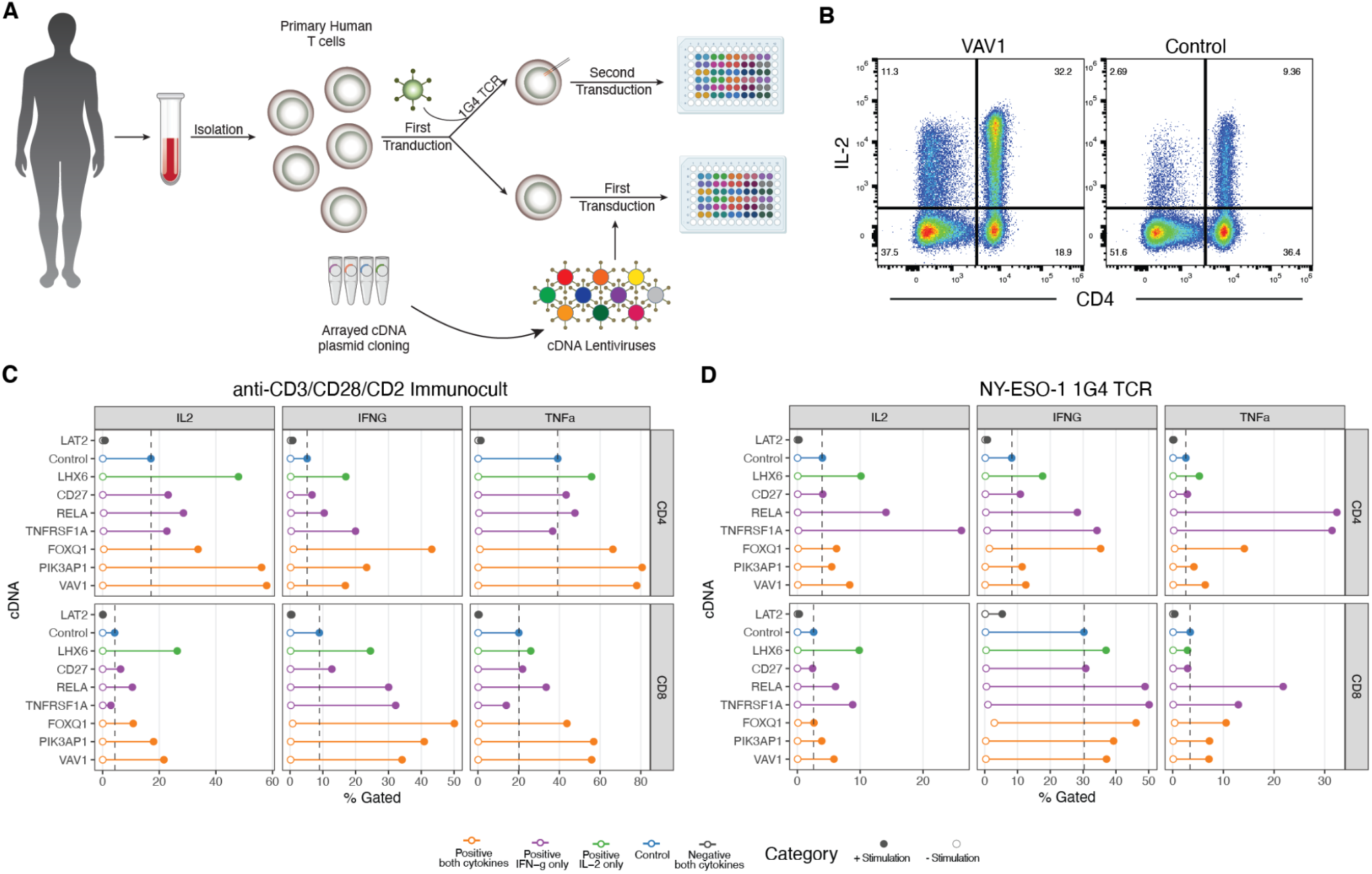
CRISPRa gain-of-function findings translate to transgenic cDNA overexpression and TCR stimulation with antigen. **(A)** Schematic of arrayed overexpression experiments with cDNA**. (B)** Flow cytometry plot for *VAV1* or Control cDNA overexpression after stimulation with Immunocult. **(C-D)** Mean Percentage of cells in cytokine positive gates from two donors after stimulation with Immunocult (C) or NALM6 target cells engineered to express NY-ESO-1 antigen in an HLA-A0201 context (D) N=2 donors.

**Figure S10.**
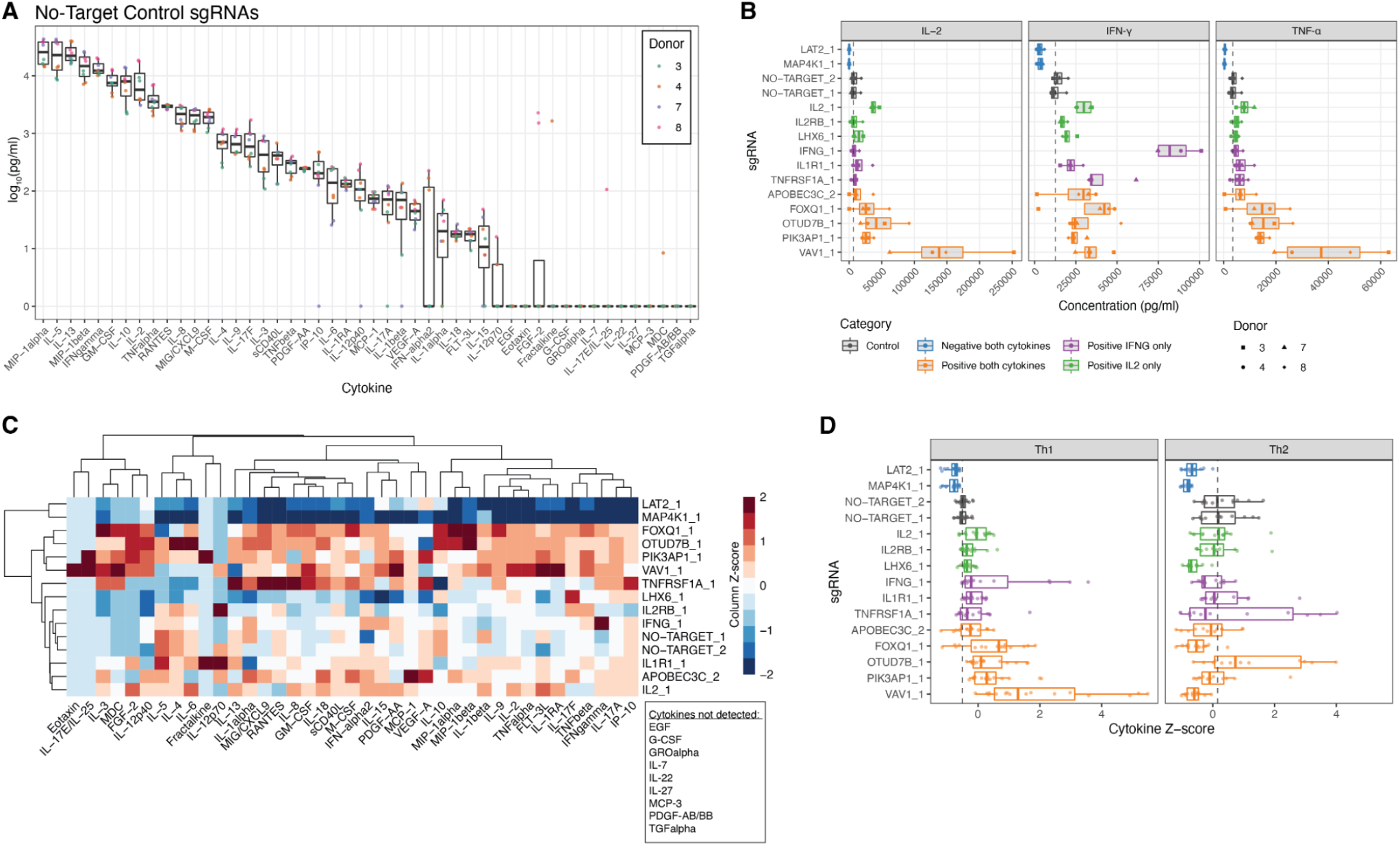
Secreted cytokine measurements in an arrayed CRISPRa panel. **(A)** Cytokine measurements in non-targeting control sgRNA samples after stimulation (2 individual sgRNAs). **(B)** Measurements of IL-2, IFN-γ, and TNF-ɑ in samples with indicated sgRNAs. **(C)** Unsupervised clustering of full cytokine panel measurements across different sgRNAs. Each tile represents the median value of 4 donors, z-score scaled across each cytokine. **(D)** Th1 and Th2 category grouped cytokine measurements across different sgRNAs. Th1 group includes of IFN-γ IL-2, TNF-ɑ, and TNF-β. Th2 group includes IL-4, IL-5, and IL-13. Each point represents a donor, cytokine measurement, z-score scaled for each cytokine.

**Figure S11.**
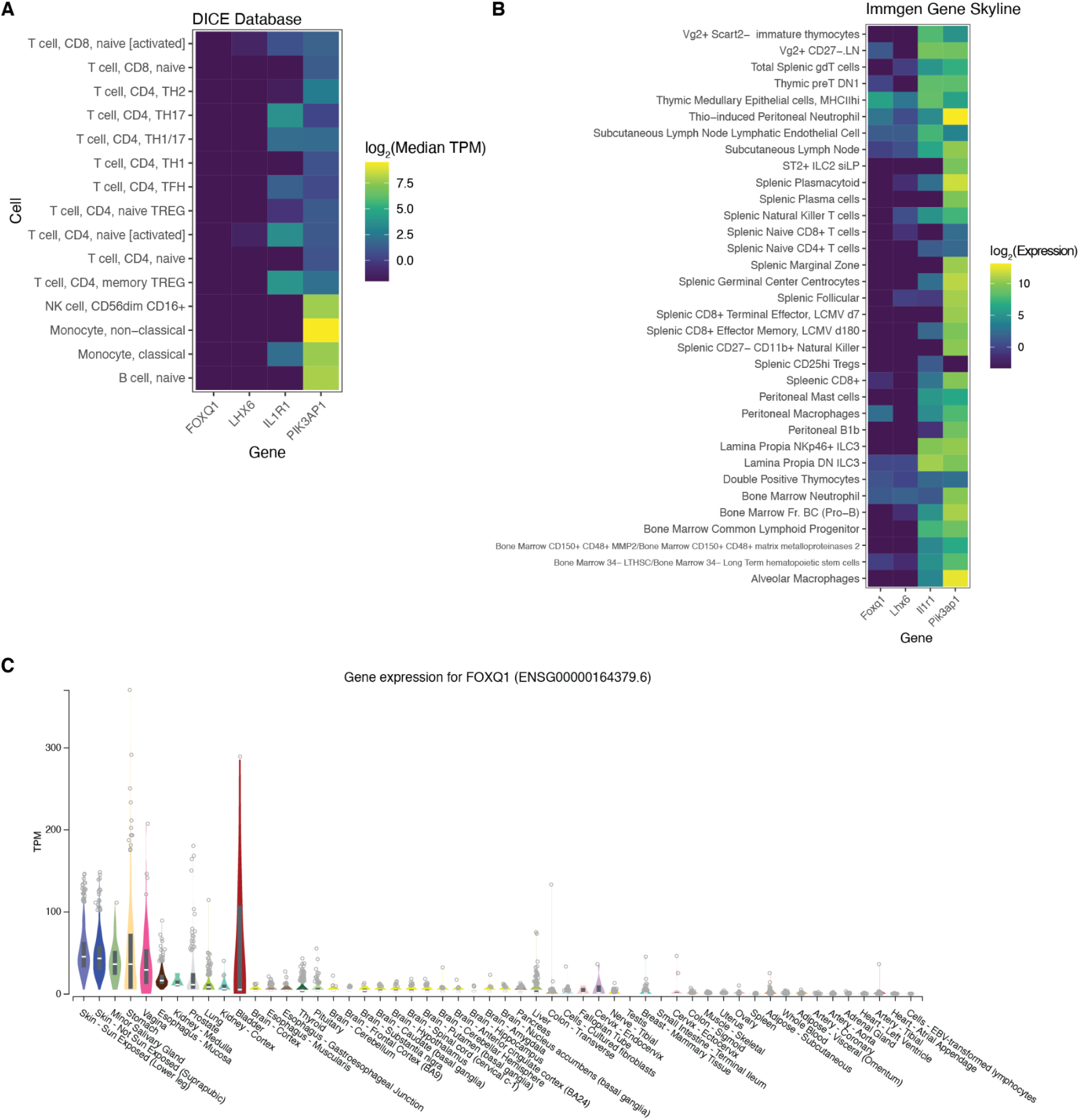
Expression of genes with low expression in T cells in publicly available transcriptome datasets. **(A)** Heatmap showing the expression of indicated genes in indicated human immune cell types, from the Database of Immune Cell Expression (DICE) (*34*). Values represent the median log_2_TPM for each gene. **(B)** Heatmap showing the expression of indicated genes in indicated mouse immune cells, from the Immunological Genome Project (Immgen) gene skyline database (*35*). **(C)** Expression of *FOXQ1* in various human tissues, from the Genotype-Tissue Expression (GTEx) project (*49*).

**Figure S12.**
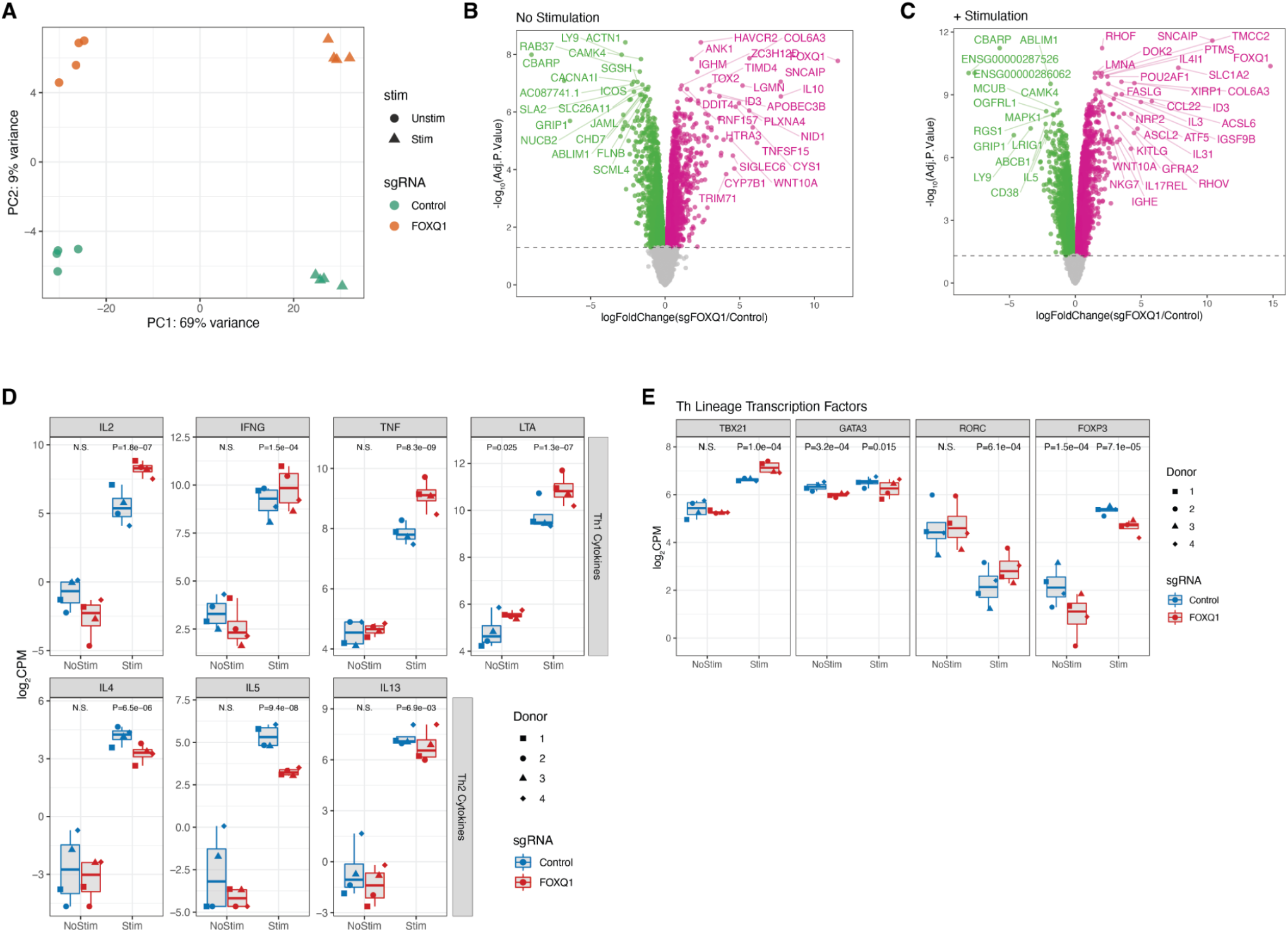
RNA-seq analysis of *FOXQ1* CRISPRa in CD4^+^ T cells. **(A)** Principal component analysis across all samples in RNA-seq transcript quantifications. **(B-C)** Differential gene expression volcano plots, between *FOXQ1* sgRNA and non-targeting control sgRNA in non-stimulated (B) and stimulated (C) T cells. **(D)** mRNA expression of Th1 signature cytokine genes (*IL2, IFNG, TNF, LTA*), and Th2 signature cytokine genes (*IL4, IL5, IL13*) across RNA-seq samples. FDR adjusted P-values between control and *FOXQ1* sgRNA samples are shown at top. **(E)** mRNA expression of T helper cell lineage defining transcription factors across RNA-seq samples. FDR adjusted P-values between control and *FOXQ1* sgRNA samples are shown at top. *TBX21* defines Th1, *GATA3* defines Th2, *RORC* defines Th17, and *FOXP3* defines Treg.

## Supplementary Tables (Available as additional excel files)

**Table S1. MAGeCK gene summaries**. Output of MAGeCK robust ranking algorithm tests from CRISPRa and CRISPRi screens.

**Table S2. Screen hits.** Gene level summary values and boolean call for screen hit in CRIPSRa and CRISPRi screens.

**Table S3. sgRNA sequences.** Sequences of single sgRNAs cloned and used in arrayed experiments in this study.

**Table S4. Flow antibodies.** List of flow cytometry antibodies used in this study.

**Table S5. RT-qPCR probes.** List of RT-qPCR probes used in this study.

